# Developmental Dysregulation of Synaptic and Myelin-Related Genes in Frontal Cortex and Serum Infrared Spectroscopy Signature in the Valproic Acid Model of Autism

**DOI:** 10.1101/2025.10.04.680465

**Authors:** Carolina Araújo, José Kroll, Juliana Alves Brandão, Sérgio Ruschi, Camilo LM Morais, Marfran Claudino Domingos dos Santos, Rafael dos Santos de Bessa, Kassio MG Lima, Sandro Souza, Rodrigo N Romcy-Pereira

## Abstract

Neural circuits emerge during development through dynamic interactions between genetic instructions and environmental cues that shape cell fate, connectivity, and the timing of myelination. In developmental disorders such as autism, abnormalities in sensory processing, social cognition, and motor behavior are thought to arise from disruptions in these processes. Here, we investigated early-life molecular changes following prenatal exposure to valproic acid (VPA), an environmentally induced model of autism. We integrated cortical gene expression analysis using RNA sequencing/qPCR, alternative splicing profiling, in situ myelin quantification, plasma serotonin determination, and machine learning-assisted classification of blood serum molecular profiles using infrared spectroscopic (FTIR) data. Our findings revealed downregulation of myelin-associated genes and upregulation of synapse-related genes in the frontal cortex of young VPA-exposed rats. qPCR confirmed reduced cortical expression of *Mobp* and *PLP1* along with increased *Penk* and *C1ql3* expression. Alternative splicing analysis identified numerous novel transcript variants, enriched in synaptic-related genes, indicating widespread post-transcriptional remodeling in VPA animals. These molecular alterations were accompanied by a significant reduction in myelin content within the cingulate and motor cortex of adult animals. Peripheral molecular profiling showed elevated plasma serotonin in VPA-treated animals and demonstrated that a support vector machine trained on serum FTIR spectra classified VPA-exposed animals with 85% accuracy. Collectively, our findings suggest that prenatal VPA exposure induces early dysregulation of myelin organization, synaptic gene networks, and RNA splicing programs, potentially leading to long-term impairments in neuronal communication and processing efficiency. Furthermore, our results highlight serum spectroscopic signatures as promising peripheral biomarkers for autism, warranting further investigation.

## 1 INTRODUCTION

During brain development, the interaction between gene expression patterns and environmental signals regulate the formation of neural circuits by coordinating neurogenesis, neuronal migration, synaptogenesis, and myelination within specific time windows (Stiles & Jernigan, 2010; Tau & Peterson, 2010). Disruptions along this sequence of events are known to affect the structural and functional organization of the nervous system, leading to physiological and behavioral abnormalities, some of them observed in autism spectrum disorder (ASD). Although genetic factors contribute substantially to ASD etiology, accumulating evidence indicates that environmental factors—such as prenatal exposure to drugs, infections, nutritional deficiencies, and pollutants— modulate the risk of the disease (Pardo & Eberhart, 2007; Modabbernia et al., 2017).

Prenatal exposure to valproic acid (VPA) is known to markedly increase the risk of ASD (Moore et al., 2000; Christianson et al., 1994; Bromley et al., 2013; Christensen et al., 2013). In animal research, the VPA model has emerged as one of the most robust and reproducible environmental models of ASD (Nicolini & Fahnestock, 2018). Rodents prenatally exposed to VPA exhibit a constellation of ASD-like phenotypes, including delays in postnatal developmental milestones, social interaction deficits, hyperlocomotion, and repetitive behaviors (Schneider & Przewłocki, 2005; Schneider et al., 2008; Kataoka et al., 2013). These animals display abnormalities reminiscent of those described in individuals with ASD, such as early brain overgrowth, reduced cerebellar volume, and a decreased number of Purkinje cells (Ingram et al., 2000; Lajeunie et al., 2001; Kim et al., 2011; Go et al., 2012; Skefos et al., 2014). Additional alterations include reduced neuronal density in the somatosensory cortex and medial prefrontal cortex (mPFC), as well as a decreased number of parvalbumin-positive (PV+) inhibitory interneurons in multiple cortical regions, including the anterior cingulate and prelimbic areas (Gogolla et al., 2009; Hara et al., 2012). Impairments in sensory processing have also been documented, such as deficits in auditory cortical responses (Anomal et al., 2015), paralleling sensory abnormalities observed in human ASD.

The clinical importance of this model is underscored by the continued widespread use of VPA, given its effectiveness and tolerability in the management of epilepsy, bipolar disorder, and migraine prophylaxis (Tomson et al., 2016). Mechanistically, VPA acts through multiple pathways. Its best-characterized molecular target is histone deacetylase (HDAC), whose inhibition alters chromatin structure and modulates gene transcription via epigenetic regulation (Phiel et al., 2001). Beyond epigenetic effects, VPA enhances GABAergic neurotransmission, blocks voltage-gated sodium channels, and influences intracellular signaling cascades that regulate neuronal differentiation and survival (Johannessen & Johannessen, 2003). These pleiotropic actions suggest that VPA exposure during critical periods of brain development may alter both the excitatory–inhibitory balance of cortical circuits and the transcriptional programs necessary for axon ensheathment and synaptic stability.

Recent transcriptomic studies in the VPA model have highlighted alterations in genes related to synaptic organization, axonal guidance, and myelin-associated proteins (Kataoka et al., 2013; Olexová et al., 2022). These findings align with neuropathological and imaging data in ASD, which point to disrupted connectivity in long-range association pathways and compromised white matter integrity, particularly within the frontal lobes (Ameis & Catani, 2015; Deoni et al., 2015). Myelination is increasingly recognized not merely as an insulating process but as a dynamic modulator of conduction velocity and spike-timing synchrony across distributed neural networks (Fields, 2015). Alterations in oligodendrocyte differentiation and myelin plasticity could therefore contribute to the atypical cortical dynamics and reduced processing efficiency characteristic of ASD (Chang et al., 2018). However, the mechanistic links between early transcriptional changes and later myelin abnormalities remain poorly understood in the VPA model.

Parallel to brain-based analyses, there is growing interest in identifying accessible peripheral biomarkers that reflect central nervous system alterations in ASD. Blood serum offers a minimally invasive source of molecular information, potentially capturing systemic correlates of neurodevelopmental dysregulation. Fourier-transform infrared (FT-IR) spectroscopy is a powerful biochemical fingerprinting technique that can detect subtle differences in the composition of proteins, lipids, nucleic acids, and carbohydrates within biofluids (Hands et al., 2014; Baker et al., 2014). Coupling FTIR data with machine learning algorithms has shown promise in classifying neurological and psychiatric conditions (Paraskevaidi et al., 2015). Yet, its application to ASD models, and particularly its integration with molecular data from brain tissue, remains largely unexplored.

In this study, we adopted an integrated approach to characterize molecular alterations in the brain and plasma of rats prenatally exposed to VPA. RNA sequencing was used to profile gene expression in the frontal cortex of infant animals, with qPCR validation of key transcripts associated with synaptic function, myelination, and VPA-specific alternative splicing. Region-specific myelin content was then assessed in adult brains using histological analyses of the corpus callosum and frontal cortical subregions. In parallel, we quantified plasma serotonin levels and applied machine learning to FTIR serum spectra to evaluate the discriminatory power of peripheral molecular profiles. Together, this approach aimed to identify early-life mechanisms linking synaptic dysregulation and myelin deficits and to explore the translational potential of spectroscopic biomarkers for ASD.

## 2 MATERIALS AND METHODS

### 2.1 ANIMAL MODEL OF AUTISM

The VPA model of autism was generated using Wistar rats (Rattus norvegicus) provided by the animal facility of the Brain Institute. All subjects were housed under standard conditions of controlled temperature of 23 ± 2°C, air-humidity 65–75%, and a 12-hour light/dark cycle with free access to food and water. Young adult female rats (P75–P80) with no prior pregnancies had their estrous cycles monitored by microscopic analysis of vaginal smears and were mated with age-matched males early in the estrus phase. Pregnancy onset was confirmed by the presence of a vaginal plug and/or spermatozoa in the vaginal smear, marking embryonic day zero (E0). At E12.5, each pregnant female received an intraperitoneal injection of sodium valproate (500 mg/kg, Sigma Aldrich) dissolved in 0.9% NaCl (250 mg/mL). Control females received an equivalent volume of 0.9% NaCl at the same gestational stage. After the injection, female behavior was observed for 30 minutes in order to ensure no side-effects besides sedation, before returning them to the animal facility. Pregnancy progression and maternal health were monitored by tracking body weight and nesting behavior until E17. Following delivery, pups remained with their mothers until the experiment day or until P21, when they were weaned and housed in same-sex cages. A total of seven VPA-exposed litters and seven control (CTL) litters were used in the experiments. All experimental procedures were approved by the Ethical Committee for Animal Use in Research of the Federal University of Rio Grande do Norte (Protocol# 0132014).

### 2.2 DEVELOPMENT AND BEHAVIORAL ASSESSMENT

#### Development

Postnatal development and prepuberal behaviors were evaluated from postnatal day (P) 1 to 35. Pregnancy was monitored by observing nesting behavior and evaluated by quantifying the number of live pups per litter. Postnatal development was assessed by observing the day pups opened their eyes and by monitoring body weight from P7 to P35.

#### Locomotor Activity

Locomotor activity was assessed by evaluating the animals’ exploratory drive in a novel environment. The task was conducted in a rectangular wooden arena (66 × 57 × 40 cm^3^) with a black floor. To encourage exploration, the walls contained evenly spaced circular holes (2 cm in diameter), arranged in two rows (six holes on the longer walls and four on the shorter walls). The apparatus was placed in a dimly lit room (illuminated by a 40W red light bulb) to minimize stress. One week before testing, all animals underwent daily handling sessions (5 minutes per day) to habituate them to the experimenter. On the test day (P30), each animal was placed in the arena and allowed to explore freely for 5 minutes (Figure 4B). Sessions were conducted between 8:00 and 10:00 AM and recorded (70 dpi HD; 30 fps; Logitech C920) for offline analysis. Locomotor activity was quantified as the total distance traveled during the session. After the test, animals were returned to their home cages and the animal facility.

#### Grooming Behavior

Grooming is an innate behavior in rodents modulated by anxiety levels (McFarlane, Kusek et al., 2008) and follows a stereotypical anteroposterior sequence, including licking of the forelegs, belly, and back. This behavior was assessed at P30 as a measure of stereotypy, a common autistic phenotype. Grooming was quantified during the locomotor activity task by measuring the frequency of self-cleaning behaviors and the average duration of each grooming event.

#### Social Interaction

Social interaction was evaluated by testing the animals’ preference for exploring an unfamiliar conspecific versus an empty cage. One week before testing, animals were habituated to the experimenter. Habituation to the experimental apparatus (P33) was conducted over two sessions (5 minutes each) in a three-chambered rectangular wooden box (105 × 60 × 35 cm^3^), divided into equal-sized compartments (35 × 60 × 35 cm^3^) connected by openings. On test day (P34), an unfamiliar rat was placed inside a metal mesh cage in one of the outermost compartments (social chamber), while an identical but empty metal cage was placed in the opposite compartment (empty chamber). The middle compartment was left empty. Each test animal was placed in the middle compartment and allowed to explore all three chambers freely for 10 minutes. The session was video recorded for offline analysis, and the time spent in each compartment was quantified. *A social discrimination index (SDI)* was calculated using the formula: *SDI = (Time in social chamber* −*Time in empty chamber)/ (Time in social chamber + Time in empty chamber)*. An SDI of 1 indicates the animal spent the entire session in the social chamber, a value close to 0 indicates equal time in both chambers, and a value of −1 indicates exclusive preference for the empty chamber. After testing, animals were returned to their home cages and transported back to the animal facility.

### 2.3 TISSUE COLLECTION AND PROCESSING

#### RNA Sequencing and qPCR

For RNA sequencing (RNA-seq) and quantitative PCR (qPCR) experiments, fresh frontal cortex tissue was carefully dissected on an ice-chilled platform. Prior to tissue collection, all surgical instruments and the workspace were sanitized with 70% ethanol and an RNase decontamination solution (RNAseZap™, Sigma Aldrich). Briefly, P15 rats were decapitated using a guillotine, and their brains were rapidly removed and washed in fresh ice-cold 0.9% NaCl. The frontal cortex from both hemispheres (+3.7 to −1.0 mm anteroposterior, referenced to bregma; Paxinos and Watson, 2008) was dissected using a surgical spatula and immediately immersed in ice-cold tubes containing RNAlater® solution. Tissue samples were kept overnight at 4°C, after which the RNAlater® solution was discarded, and the samples were stored at −80°C until further processing.

#### Histological Processing for Myelin Analysis

For myelin histological experiments, brain tissue was processed following transcardial perfusion at P15 and P60. Animals were anesthetized with sodium thiopental (80 mg/kg, i.p.) and perfused with room-temperature phosphate-buffered saline (PBS; 10 mM), followed by cold 4% paraformaldehyde (PFA) in phosphate buffer (PB; 100 mM). After removal from the skull, brains were post-fixed in 4% PFA overnight at 4°C. The next day, brains were transferred to a 30% sucrose solution in PBS (10 mM) until fully submerged. Brains were then rapidly frozen in dry ice-chilled isopentane and stored at −80°C until cryostat sectioning (Microm HM 550). Coronal sections (40 µm) were collected in cell culture plates and stored free-floating in a cryoprotectant solution (0.1 M PB, 30% glycerol, and 30% ethylene glycol) at −20°C until further processing. Tissue sections were mounted on slides prior to staining.

#### Serotonin quantification and infra-red spectroscopy of blood plasma

Blood samples were collected in plastic tubes and stored at −80°C. On experiment day, samples were centrifuged (2000g, 10 min at 4°C) to separate the cellular content from the supernatant (blood plasma). Plasma samples were used for the HPLC and FTIR analysis. Plasma serotonin quantification was conducted in individual samples from P15 pups. Infrared spectroscopy was performed on samples collected from infant, adolescent, and adult animals spanning postnatal days 8 to 60 (P8– P60), with individual spectra recorded for each sample.

### 2.4 RNA SEQUENCING AND ANALYSIS

#### RNA sequencing

Total RNA from frontal cortex tissue was extracted and purified using a monophasic solution of phenol and guanidinium isothiocyanate, the Trizol method (TRI Reagent® Solution, Ambion, Carlsbad, CA, USA). RNA samples from VPA (n = 6 animals) and CTL (n = 5 animals) groups were pooled together in tubes for each group, and then precipitated using 3M Sodium Acetate and 100% Ethanol. Before sequencing, RNA concentration and quality were checked by Qubit Fluorometer (Thermo Fisher Scientific, Waltham, MA, USA) and 2100 Bioanalyzer (Agilent, Santa Clara, CA, USA) using Agilent RNA 6000 nano reagents, respectively. RNAseq libraries were constructed with the purified RNA samples (RIN scores 8.6-8.7). RNA sequencing was performed using HiSeq™ 2000 Next-Generation Sequencing platform (Illumina, San Diego, CA, USA) and was run as a service by BGI (http://genomics.cn). Briefly, this platform runs automated clonal amplification for cluster generation, and uses the reversible terminator sequencing-by-synthesis method for RNA sequencing.

#### Alignment of reads

The rat genome reference sequence (RGSC/rn5) was downloaded from UCSC Genome Bioinformatics portal (http://genome.ucsc.edu). Transcriptomic reference data was downloaded from Ensembl, release 76 (ftp://ftp.ensembl.org/pub/release-76/gtf/rattus_norvegicus). RNA-Seq data from CTL and VPA samples were mapped to the reference genome using Tophat version 2 (Trapnell et al., 2009). A conservative parameter was maintained (--splice-mismatches 0), and a reference GTF file, containing transcriptomic alignments from Ensembl, was set for helping the mapping process.

#### Differential Expression Analysis

RNA-Seq data from CTL and VPA rats were mapped to the reference genome using Tophat version 2. A conservative parameter was maintained (--splice-mismatches 0), and a reference GTF file, containing transcriptomic alignments from Ensembl, was set for helping the mapping process. The resulting BAM files from Tophat were used as input for HTSeq-count (HTSeq, 2014), which returned the raw number of reads for each gene. Afterwards, gene counts from CTL and VPA samples were submitted to DESeq (Anders and Huber, 2010) R-package for a differential expression analysis based on a model using the negative binomial distribution.

#### Gene Ontology and Functional Interpretation

Gene ontology enrichment and functional analysis of RNAseq differentially expressed genes was carried out with help of clusterProfiler (Yu et al., 2012) package for R and web applications Panther v.14 (Gene Ontology Consortium), KEGGS pathway, QuiGO/EMBL-EBI and SFARI gene database Release 2025-Q2 (Ashburner et al., 2000; Gene Ontology Resource, 2021; Simon Foundation Autism Research Initiative). Statistical significance was calculated by Fisher test followed by Benjamini-Hochberg method (false discovery rate, FDR). Significance was considered for adjusted p-values at FDR level of 0.05.

#### Chromosomal mapping of differentially expressed genes

We searched the location of each VPA-related DEG in the rat and human chromosomes using computational tools available at the Rat Genome Database-RGD (Vedi et al., 2023) and cross-checking in the NCBI, UCSC genome browser and EMBL-EBI. Rat genome assembly Rnor_6.0 and human genome assembly GCh38 were used.

#### Alternative Splicing Analysis

The resulting BAM files created by Tophat were used as input to Cufflinks version 2 software, using the Ensembl reference data, in order to obtain the transcriptome assembly (Roberts et al., 2011). Thereafter, alternative splicing analysis was performed as previously described by our group (Galante et al., 2004; Galante et al., 2007; Kroll et al., 2012). Briefly, the genome coordinates of all expressed sequences were compared to the genome coordinates of Reference Sequence (RefSeqs) entries. Only sequences with multiple exons and sharing at least one exon-intron boundary with a RefSeq sequence were merged. The sequences were then annotated with gene symbols, complying with the HUGO (Gene Nomenclature Committee) standards. Alternative splicing events (ASE) were identified by a pairwise comparison algorithm, which is a common approach in other similar tools (Florea et al., 2012; Kroll et al., 2012; Sammeth et al., 2008). Furthermore, we used a computational strategy previously implemented in Splooce (Kroll et al., 2012), which is based on regular expressions capable of identifying all well-known simple and combined ASEs, such as exon skipping, intron retention and alternative 5’ and 3’ splicing borders (Kroll et al., 2015). Therefore, we assigned each ASE to a one class, quantified their expression levels and mapped the alternative spliced transcripts to RefSeq. Splicing variants were classified as known and unknown based on their presence or absence in the RefSeq database annotation, respectively. VPA-specific variants were analyzed considering transcripts with >50 FPKM and relative expression >75% than CTL.

### 2.5 qPCR validation

Real-time quantitative PCR (qPCR) analysis was carried out in RNAseq samples (CTL, 4-5 animals; VPA, 4-5 animals) and new independent samples (control, 6-8 animals; VPA, 6-14 animals). cDNA samples were prepared as follows: Total RNA was isolated using Trizol method (TRI Reagent® Solution, Ambion, Carlsbad, CA, USA) and cDNA was synthesized from RNA by RT-PCR. Before cDNA synthesis, we treated RNA samples with DNAse I (AM2224, Thermo Fisher Scientific) and RNAse inhibitor (RNaseOUT™ Recombinant Ribonuclease Inhibitor, 10777019, Thermo Fisher Scientific) at 37°C for 30 minutes. To inactivate DNAse activity, 10mM EDTA was added to the samples for 10 minutes at 65°C. cDNA was synthesized by reverse transcription polymerase chain reaction (RT-PCR) using random oligo primers following manufacturer’s instructions (High-Capacity cDNA Reverse Transcription Kit, 4368814, Thermo Fisher Scientific). qPCR reactions were carried out in triplicates for each sample in a 96-well plate using SYBR green (SYBR® Select Master Mix, Applied Biosystems, Carlsbad, CA). Reactions were run in Viia 7 Real-Time PCR System (Applied Biosystems). Each qPCR reaction plate included a negative control (without cDNA sample), negative RT controls (RNA sample treated with DNAse, RNA inhibitor, EDTA and all the qPCR reagents, but no reverse transcriptase, thus not converted into cDNA) to check for genomic DNA contamination. Additionally, a melting curve was performed at the end of each qPCR experiment to evaluate primers’ specificity. We analyzed gene expression using the relative quantification method – ΔΔCt, cycle threshold (Ct), where fold change is equal to 2^-ΔΔCt^ (Livak and Schmittgen, 2001). Before the experiments, we tested the amplification efficiency of each primer compared to the endogenous control (HPRT) in PCR reactions on five serial dilutions of a control cDNA sample. Cycle threshold (Ct) at each dilution was plotted as a standard curve for each primer. Efficiency was calculated according to the manufacturer (Applied Biosystems) and only primers with values between 90% and 100% were used. The following primers (5’-3’) were used: Mobp (product size: 91 bp), *forward*: tgaaaacacagtaagatgagtcaaaaag (Tm = 61.2°C), *reverse*: agtggatgctgaagtgctctga (Tm = 62.5°C); Mag (product size: 105 bp), *forward*: tggaagcccacagtgaat (Tm = 57.5°C), reverse: cttgaagatggtgagaataggg (Tm = 57.8°C); Plp1 (product size: 137 bp), forward: tgcgctgatgccagaatgt (Tm = 64.0°C), *reverse*: gaaggttg-gagccacaaacc (Tm = 61.8°C); Klhl1 (product size: 123 bp), *forward*: ggccagtgatgacgtaaat (Tm = 56.0°C), reverse: agtcttatgaaggcgaggag (Tm = 56.2°C); Penk (product size: 199 bp), *forward*: cgagttcccttgggataaca (Tm = 59.9°C), reverse: catgaaaccgccatacctct (Tm = 60.0°C); C1ql3 (product size: 120 bp), *forward*: acgaggtgctcaagttcg (Tm = 57.4°C), reverse: gcatcaggacgtggtagg (Tm = 57.5°C); HPRT (product size: 146 bp), forward: ggccagtaaagaactagcaga (Tm = 56.4°C), *reverse*: cctacaggctcatagtgcaa (Tm = 56.6°C).

### 2.6 MYELIN STAINING

Myelin staining was performed in brains slices of VPA and CTL animals. Prior to staining, brain sections were re-hydrated in 10 mM PBS for 10 minutes and then immersed in 0.3% Black-Gold II staining solution (Black-Gold II, Histo-Chem Inc.) diluted in 0.9% saline, for 15 minutes at 60°C. Slices were then washed in 10 mM PBS (3 times, 5 minutes each) and fixed with 1% sodium thiosulfate solution (Sigma Aldrich) for 3 minutes at 60°C. After three additional washes in 10 mM PBS (5 minutes each), sections were dehydrated consecutively in 50%, 70%, 95%, and 100% ethanol solutions for 3 minutes each, followed by 30 minute or longer incubation in xylene until mounting with DPX (44581, Sigma Aldrich). Staining was done in several batches composed of paired VPA and CTL slices.

To quantify the intensity of myelin staining in the cortex and corpus callosum, we captured 20X brightfield images and stitched them together (mosaic together) using a Zeiss Imager M.2 ApoTome 2 microscope (Carl Zeiss MicroImaging, Thornwood, NY). All slides were coded and blind analysis was performed using Fiji software (Schindelin et al., 2012). Before analysis, mosaics were converted to 8-bit images and regions of interest (cortex and corpus callosum) were delimited based on the Atlas of the Developing Rat Brain in Stereotaxic Coordinates P14 (Khazipov et al., 2015) for the coronal slices of P15 animals; and on The Rat Brain in Stereotaxic Coordinates (Paxinos and Watson, 2008) for coronal slices of P60 animals, and for sagittal slices of P15 and P60 animals.

Myelin staining in frontal cortex slices was quantified in the coronal plane from the left hemisphere. The whole frontal cortex area was divided into five regions: infralimbic, prelimbic, cingulate, motor, and somatosensory, from +1.4 to +0.4 mm relative to bregma (Khazipov et al., 2015) for P15 animals, and from +3.7 to +0.96 mm relative to bregma (Paxinos and Watson, 2008) for P60 animals. In P15, we quantified staining intensity in 11 brain sections from 4 CTL animals (3 litters) and 9 brain sections from 3 VPA animals (2 litters). In P60, we quantified staining intensity in 16 brain sections from 4 CTL animals (2 litters) and 23 brain sections from 4 VPA animals (3 litters). Corpus callosum area was measured in the sagittal plane from the right hemisphere and ranged from 4.2 to 1.9 mm relative to the midline (Paxinos and Watson, 2008) for P15 and P60 animals. Myelin staining intensity was quantified by averaging pixel intensity in the region of interest (mean gray value). Myelin staining intensity of the frontal cortex and corpus callosum was normalized by the slide background, and corpus callosum area was normalized by the total brain area.

### 2.7 PLASMA SEROTONIN

Plasma serotonin (5-HT) levels were quantified from VPA and CTL animals at age P15. A fraction of the plasma volume was kept aside for protein quantification (Bradford, 1976). To the remaining sample we added 0.1 M perchloric acid containing 43.8 ng/mL of 3,4-dihydroxybenzylamine in order to precipitate proteins and serve as internal standard, respectively. The mix was twice centrifuged at 12,000 g for 10 minutes, 4 ºC and the supernatants collected before chemical analysis. The chromatographic analysis was conducted in an Agilent 1260 LC system (Agilent Technologies, Mississauga, ON, Canada), equipped with ANTEC electrochemical amperometric detector (ANTEC Analytical Technology, Leyden, Netherlands). Separation was performed using a Poroshell 120 EC-C18 column (50 mm × 3.0 mm, 2.7 μm, Agilent Technologies) using as mobile phase NaH_2_PO_4_·H_2_O (100 mM), Na_2_EDTA (0.1 mM) and 1-octanesulfonic acid sodium salt (0.4 mM) buffer in ultrapure water (18.2 MΩ.cm). The pH was adjusted to 3.3 with phosphoric acid. Mobile phase flow rate was set at 0.5 mL/min, temperature at 40°C (separation and detection) and detector electric potential was set as +400 mV vs. Ag/AgCl reference electrode. Samples were injected in a volume of 8 μL. Chromatography data were plotted using Open Lab™ software (Agilent Technologies, Mississauga, ON, Canada). All experimental and control samples were run in the same assay.

### 2.8 FTIR SPECTROSCOPY AND CHEMOMETRIC METHODS

A Shimadzu FTIR model IRAffinity-1, with an attached ATR accessory was used for spectral acquisition. Air was used as a background for every new sample. The spectra were recorded in the range of 4000^-1^ to 400 cm^-1^, with a resolution of 4 cm^-1^, 32 scans per spectrum, and with a time of 26 seconds to obtain each spectrum. The room temperature was 22ºC, and the samples were allowed to come into equilibrium with that temperature. Ten microliters (10 μL) of blood plasma was placed on the ATR crystal for spectral measurement. After obtaining each spectrum, the crystal was washed with 70% alcohol (v/v) and dried with paper towels. After cleaning the crystal, a new background was made before the acquisition of the next spectrum.

The spectral data processing was performed within MATLAB R2012b software (MathWorks Inc., USA) using the PLS Toolbox version 7.9.3 (Eigenvector Research Inc., USA) for pre-processing and the Classification Toolbox for MATLAB (Ballabio and Consonni, 2013) for classification. As pre-processing, the raw spectral data was cropped between 1178–2139 cm^-1^ and 2432–4000 cm^-1^ in order to remove CO_2_ interference and mere background signal present in the remaining spectral regions. Savitzky-Golay (SG) smoothing (window of 15 points, 2^nd^ order polynomial fitting) followed by automatic weighted least squares (AWLS) baseline correction (3^rd^ order polynomial fitting) were applied to the cut data in order to correct for random spectral noise and baseline distortions.

The pre-processed spectral data, derived from 30 CTL (17 males/13 females; ages P8-15: 03; P16-35: 11; P36-60: 16) and 26 VPA animals (15 males/11 females; ages P8-15: 07; P16-35: 12; P36-60: 07) animals with 3 spectra per animal, was split into training (*n* = 60 CTL, *n* = 52 VPA samples) and test (*n* = 30 CTL, *n* = 26 VPA samples) sets using the MLM algorithm (Morais et al., 2019b). The training set was used for model construction and optimization, and the test for final model evaluation. Classification was performed using a support vector machines (SVM) classifier. SVM is a binary linear classifier with a non-linear step called the kernel transformation (Cortes and Vapnik, 1995). SVM works in an attempt to define a decision limit, or hyperplane, capable of separating two classes optimally, maximizing their distances. SVM classifies data from different classes by determining a set of support vectors, through the training set. In the optimization stage, SVM seeks a decision limit with a greater margin in relation to the points closest to the two classes. To find this hyperplane that best differentiate the samples from different classes, SVM uses a kernel function. The radial basis function (RBF) is the most common kernel function, since it tends to adapt well to different data distributions (Morais et al., 2019a).The RBF Kernel is calculated as follows (Morais et al., 2017):

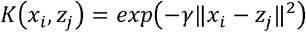

where *x* and *z* are sample measurement vectors and is the parameter that will determine the RBF width. The SVM classifier takes the form (Morais et al., 2017):

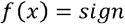

with representing the number of support vectors; *i* represents the Lagrange multipliers; *y*_*i*_ represents the class membership (± 1); *K*(*x*_*i*_,*z*_*j*_) is the kernel function; and *b* is the bias parameter. The SVM parameters were optimized using a venetian blinds cross-validation with 5 data splits in the training set, where the optimum performance was obtained with a kernel parameter of 0.2, cost of 1000, and 73 support vectors.

The classification results of the SVM model applied to the test set were validated based on some quality metrics, such as accuracy (total number of samples correctly classified considering true and false negatives), sensitivity (proportion of positives that are correctly identified) and specificity (proportion of negatives that are correctly identified), which were calculated as follows (Morais et al., 2019b):

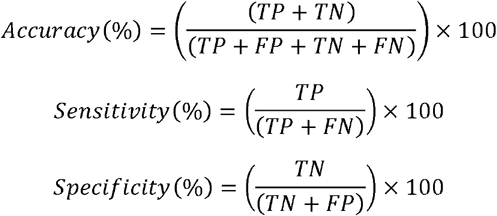

where TP stands for true positives, TN for true negatives, FP for false positives, and FN for false negatives.

### 2.9 STATISTICS

For statistical analysis, we used Student’s t-Test or Mann-Whitney U test to compare data from VPA and CTL groups under normality or not. ANOVA with repeated measures was used to compare group weights at different ages and Log-rank test to compare eye-opening curves. Statistical significance level was set to 0.05.

## 3 RESULTS

### DEVELOPMENT AND BEHAVIOR

Prenatal development and postnatal phenotypes were assessed in VPA-treated animals by measuring litter size at birth, monitoring weight gain, and performing behavioral tests from P1 to P35. Litter size did not differ between VPA-treated and control dams, indicating normal pregnancy outcomes. In contrast, VPA-exposed pups exhibited delayed eye opening and reduced post-weaning body weight (Fig. 1A–D). Juveniles (P30–P35) displayed increased locomotion and persistent grooming in the open field, indicative of heightened perseverative behavior (Fig. 1E–G). In the social interaction test, VPA animals showed no preference for a non-familiar conspecific over an empty chamber, whereas controls preferred the social chamber (Fig. 1H).

**Fig. 1.**
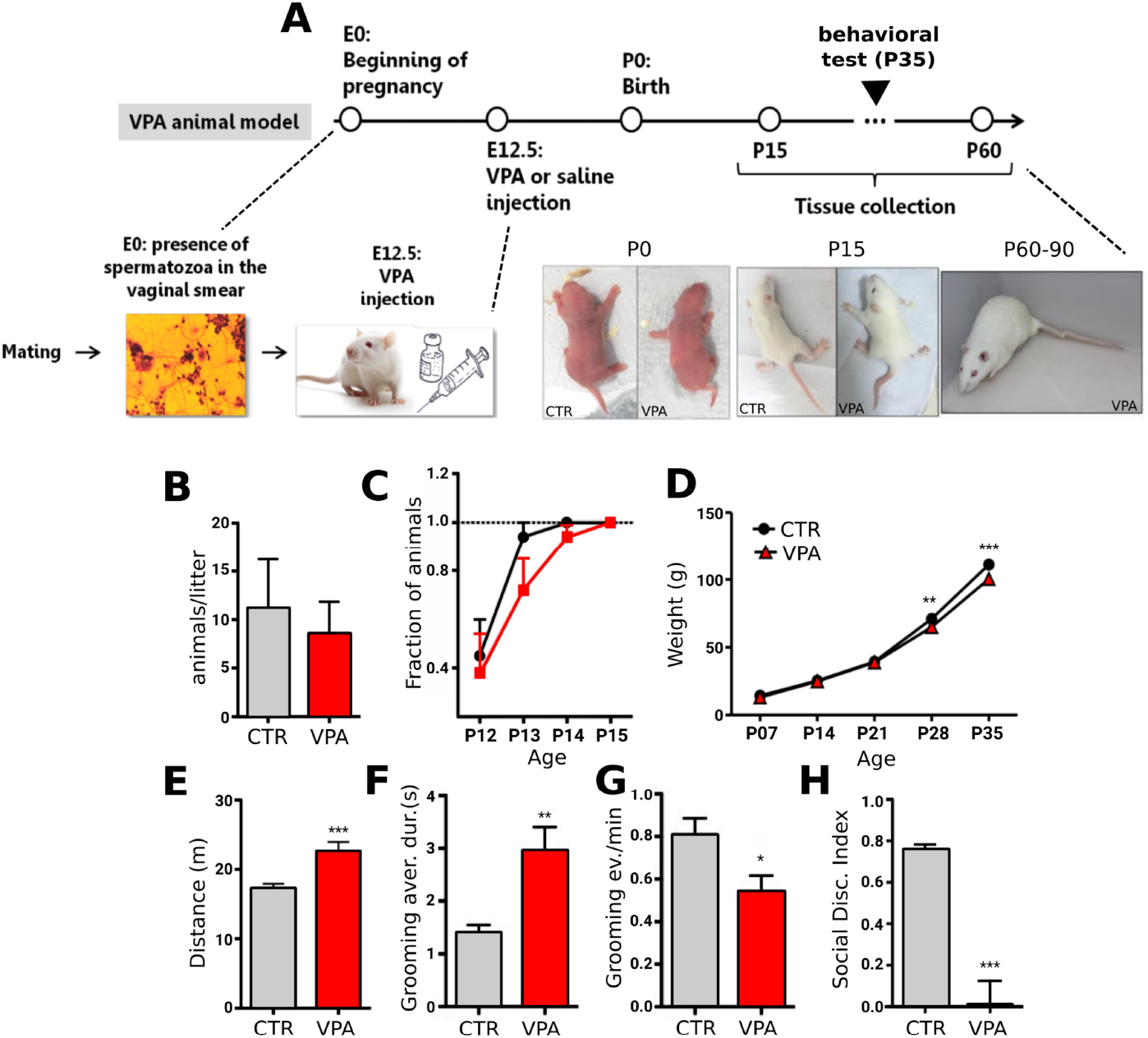

### DIFFERENTIAL EXPRESSION AND GENE ONTOLOGY ANALYSIS

Gene expression in the frontal cortex was analyzed in P15 male VPA-treated and control animals (Fig. 2). Of 16,032 genes detected, 179 were differentially expressed (DEGs) in VPA animals (p < 0.05), including 15 at p < 0.01. Among DEGs, 103 were downregulated and 76 upregulated (Fig. 2A). Expression levels ranged from 0.9 to 626.0 FPKM for downregulated genes (median = 16.5; IQR = 40.9) and 1.1 to 118.3 FPKM for upregulated genes (median = 12.7; IQR = 20.3). Fold changes ranged from −38.4% to −23.7% (median = −28.8%; IQR = 3.9%) for downregulated genes and from +31.4% to +62.8% (median = +42.3%; IQR = 7.5%) for upregulated genes (Fig. 2B–C).

**Fig. 2.**
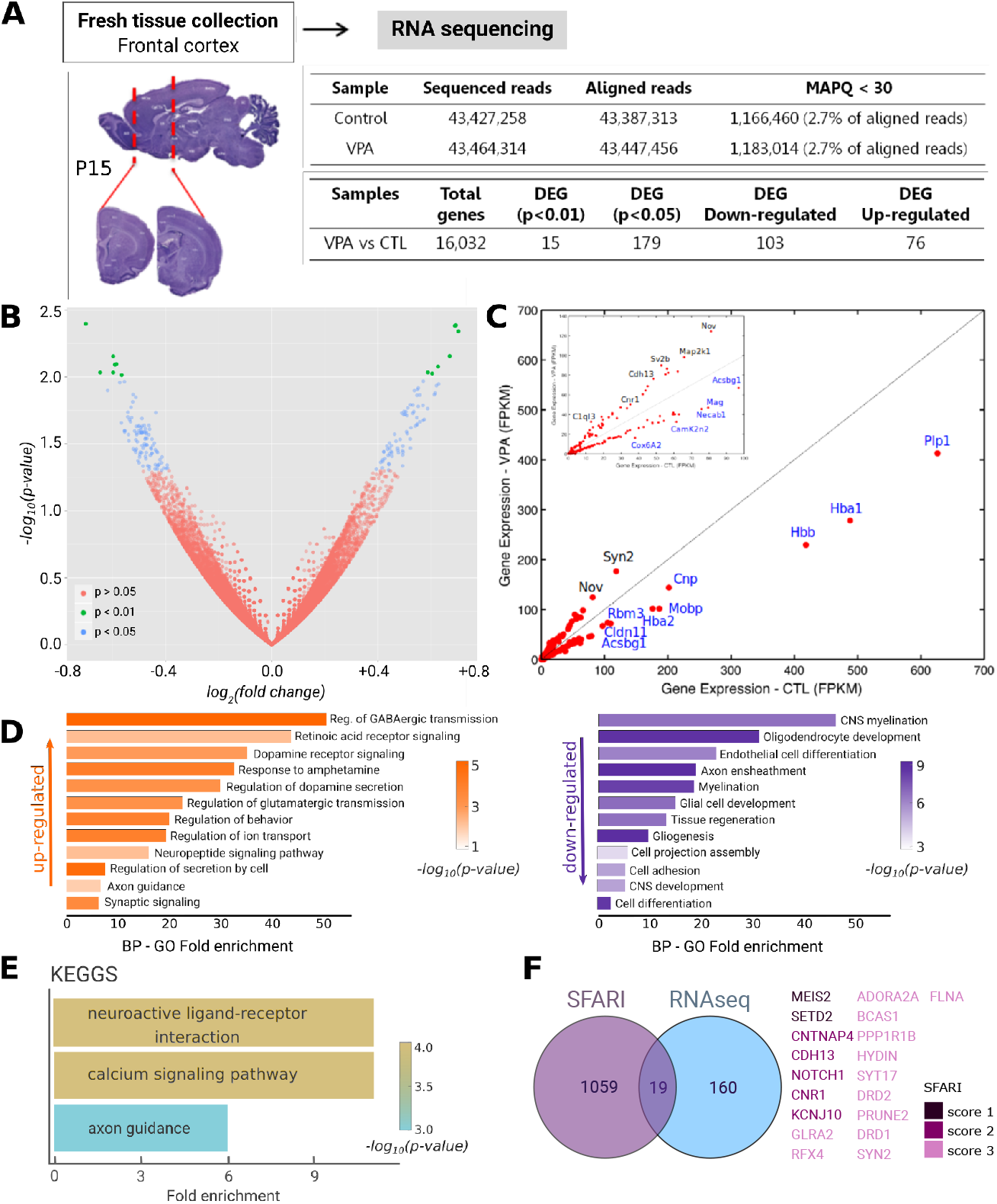

Gene ontology analysis (Fig. 2D) revealed that upregulated DEGs were enriched in synaptic signaling pathways, including GABAergic, glutamatergic, dopaminergic, and neuropeptidergic transmission: regulation of GABAergic synaptic transmission (50×; p < 2.0 × 10^−5^), regulation of dopamine secretion (30×; p < 1.0 × 10^−3^), dopamine receptor signaling (35×; p < 1.0 × 10^−3^), regulation of glutamatergic transmission (>20×; p < 2.0 × 10^−4^), neuropeptide signaling (>15×; p < 1.0 × 10^−2^), and general synaptic signaling (>5×; p < 2.0 × 10^−4^). Representative genes included Rxrg, C1ql3, Penk, Gnal, and Rgs9 (2.0–32.7 FPKM).

Downregulated DEGs were enriched in myelination-related processes: CNS myelination (>40×; p < 5.0 × 10^−7^), oligodendrocyte development (>30×; p < 6.0 × 10^−9^), endothelial cell differentiation (>20×; p < 1.0 × 10^−4^), axon ensheathment (18×; p < 2.0 × 10−8), myelination (17×; p < 5.0 × 10^−7^), and glial cell development (15×; p < 1.0 × 10^−5^). Myelin-associated genes with robust expression (>30 FPKM) that were downregulated in VPA animals included *Plp1, Cnp, Mobp, Cldn11, Mag, and Ugt8*. KEGG pathway enrichment highlighted neuroactive ligand–receptor interactions, calcium signaling, and axon guidance (Fig. 2E).

Approximately 10% of DEGs (19/179) were listed in the SFARI gene database (2025 Q2-release) as autism-associated (Fig. 2F): two high-confidence genes (score 1; *Meis2, Setd2*), five strong candidates (score 2; *Cntnap4, Cdh13, Unc79, Cnr1, Kcnj10*), and eleven with suggestive *evidence* (score 3; *Glra2, Rpl10, Adora2a, Bcas1, Ppp1r1b, Hydin, Syt17, Drd2, Prune2, Drd1, Syn2*). SFARI categories are defined as follows: S = syndromic; score 1 = high confidence (FDR < 0.1); score 2 = strong candidate; score 3 = suggestive evidence (Satterstrom et al., 2020).

Considering the list of genes implicated in autism susceptibility hosted in the SFARI database, we mapped SFARI genes to the human chromosomes and then, mapped VPA-related DEGs on rat chromosomes in order to compare their distributions (Fig. 3). SFARI assigns every gene in the database with a score reflecting the strength of the evidence linking it to the development of autism. Score 1 genes are considered of high confidence clearly implicated in autism typically by the presence of at least three de novo likely-gene-disrupting mutations, while score 2 genes are considered strong candidates with two de novo mutations uniquely implicated by a genome-wide association study consistently replicated and accompanied by evidence that the risk variant has a functional effect (SRAFI database, 2025). Score 1-2 genes were mapped to their locations in the human chromosomes, which showed that chromosomes 2, 1, X, 7, 3, 11, 5, 17, 6, 15, 19, 12 and 16 concentrated a significant fraction of them (Fig. 3A). In addition, chromosomes 2, 3, 5, 7, 9, 15 and X showed an enrichment of SFARI genes as compared to chance levels (random distribution related to the number of genes in each chromosome). Using the synteny map between the human and rat genomes, we showed that the six human chromosomes most enriched for autism-related genes mapped onto ten rat chromosomes: 1, 2, 3, 4, 6, 8, 9, 11, 18 and X (Fig. 3B). In order to compare with the chromosomal distribution of the DEGs induced by VPA, we mapped these genes to the rat chromosomes. We observed they were not randomly distributed, being particularly enriched in nine of the rat chromosomes: 19, 2, 3, 11, 13, X, 15, 8, and 4 (ordered from low-to-high enrichment; Fig. 3C). Up- and down-regulated DEGs also showed to be unevenly distributed across the rat chromosomes (Fig. 3C).

**Fig. 3.**
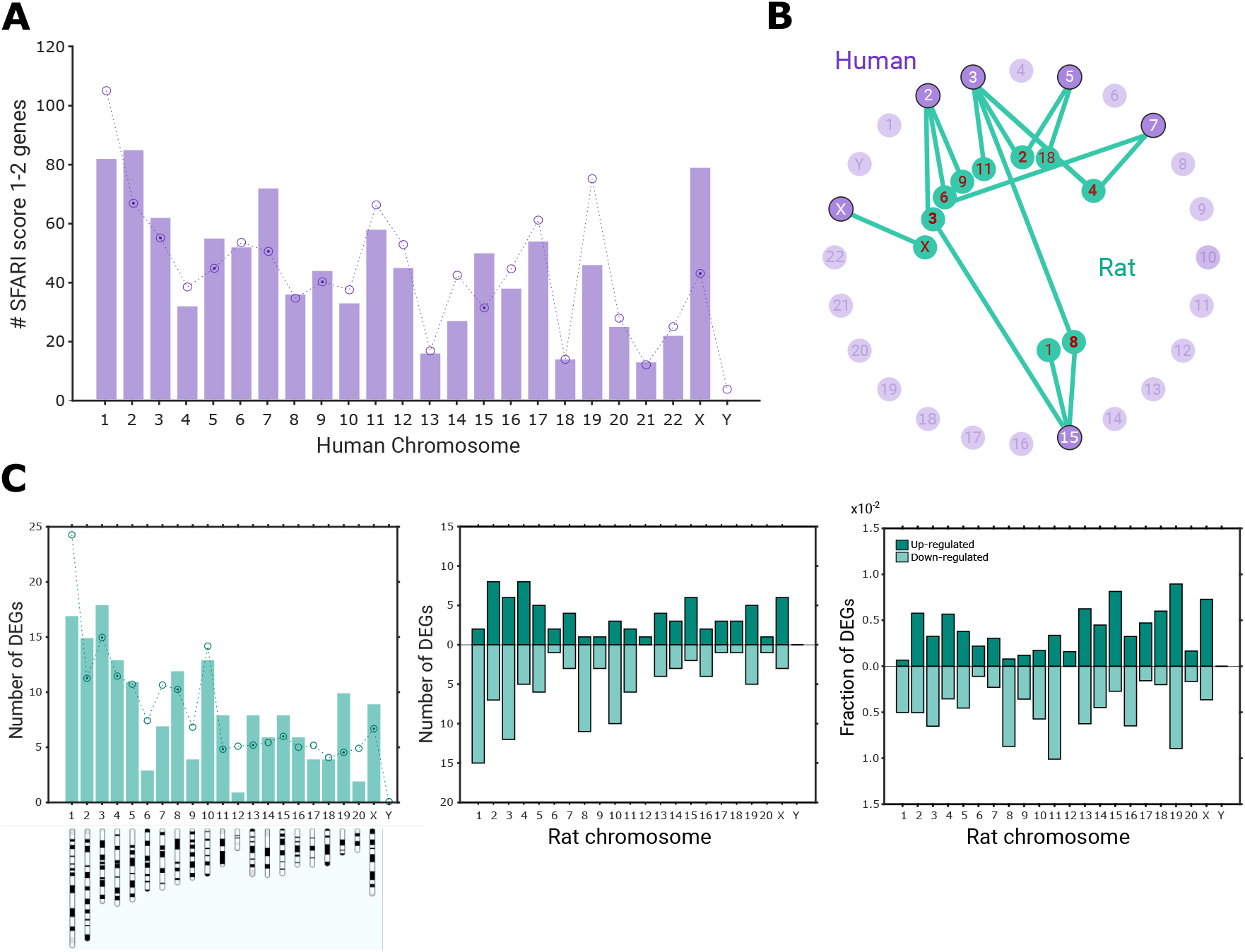

### VALIDATION OF DIFFERENTIAL EXPRESSION PROFILE

Based on the list of differentially expressed genes and the GO enrichment analysis (Fig. 2D), we selected six genes for validation by qRT-PCR. Four DEGs implicated in myelin organization during development: *Klhl1* (Kelch Like Family Member 1), *Mag* (myelin-associated glycoprotein), *Mobp* (myelin-associated oligodendrocytic basic protein), and *Plp1* (proteolipid protein 1) and two DEGs associated with synaptic transmission: *Penk* (proenkephalin; associated to dopaminergic pathways) and *C1ql3* (Complement 1q-like 3, implicated in synaptic pruning; Fig. 4A). Validation was performed using newly generated independent samples from pups of distinct litters (3, CTL; 4, VPA), as well as the original RNA-seq samples.

Our results showed that *Mobp* and *Plp1*, both critical for developmental myelination, were significantly downregulated in the frontal cortex of P15 VPA-treated animals (p < 0.05, Student’s t-test; Fig. 4B). *Klhl1* and *Mag*, in contrast, were similarly downregulated in RNA-seq samples and sequencing results, but did not differ between VPA-treated animals and controls, in the newly generated independent samples. Moreover, the enkephalin precursor, *Penk* also involved in synaptic plasticity and activity-dependent circuit refinement during development and *C1ql3*, a synaptic organizer involved in excitatory synapse formation and pruning, were upregulated in the frontal cortex of VPA animals (p < 0.05, Student’s t-test). Therefore, these transcriptional changes point to concurrent alterations in myelin-related processes and excitatory synaptic organization during early postnatal cortical development in VPA-exposed rats.

**Fig. 4.**
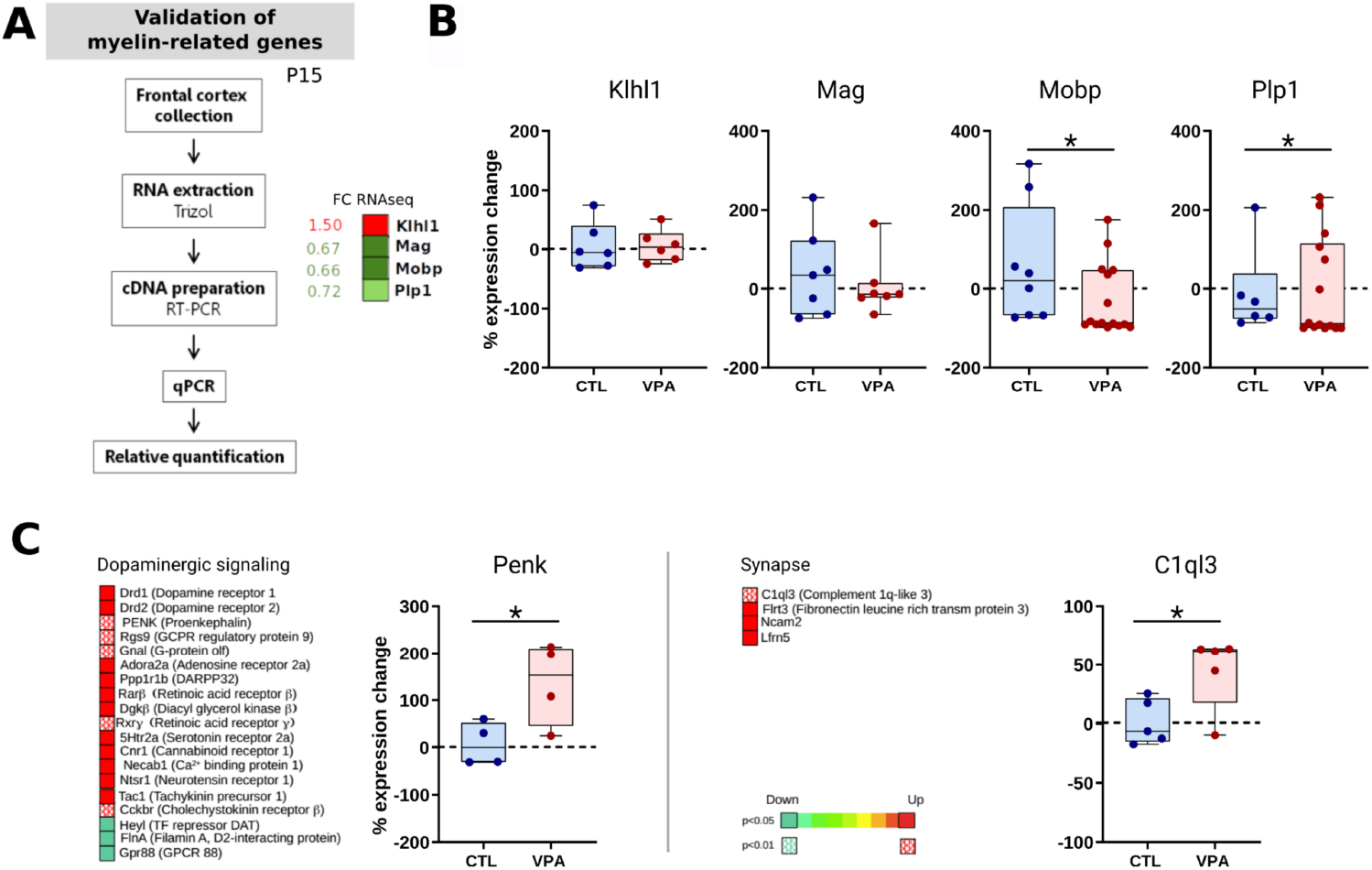

### ABNORMAL IN SITU MYELINATION

Given the transcriptional downregulation of *Mag* and *Mobp*, we next examined white matter structure and organization in VPA- and control-treated brains using Black-Gold II myelin staining, which selectively labels densely myelinated axons (Schmued et al., 2008; Mengler et al., 2014). Coronal and sagittal sections containing the frontal cortex and corpus callosum were analyzed at P15 and P60 (Fig. 5).

**Fig. 5.**
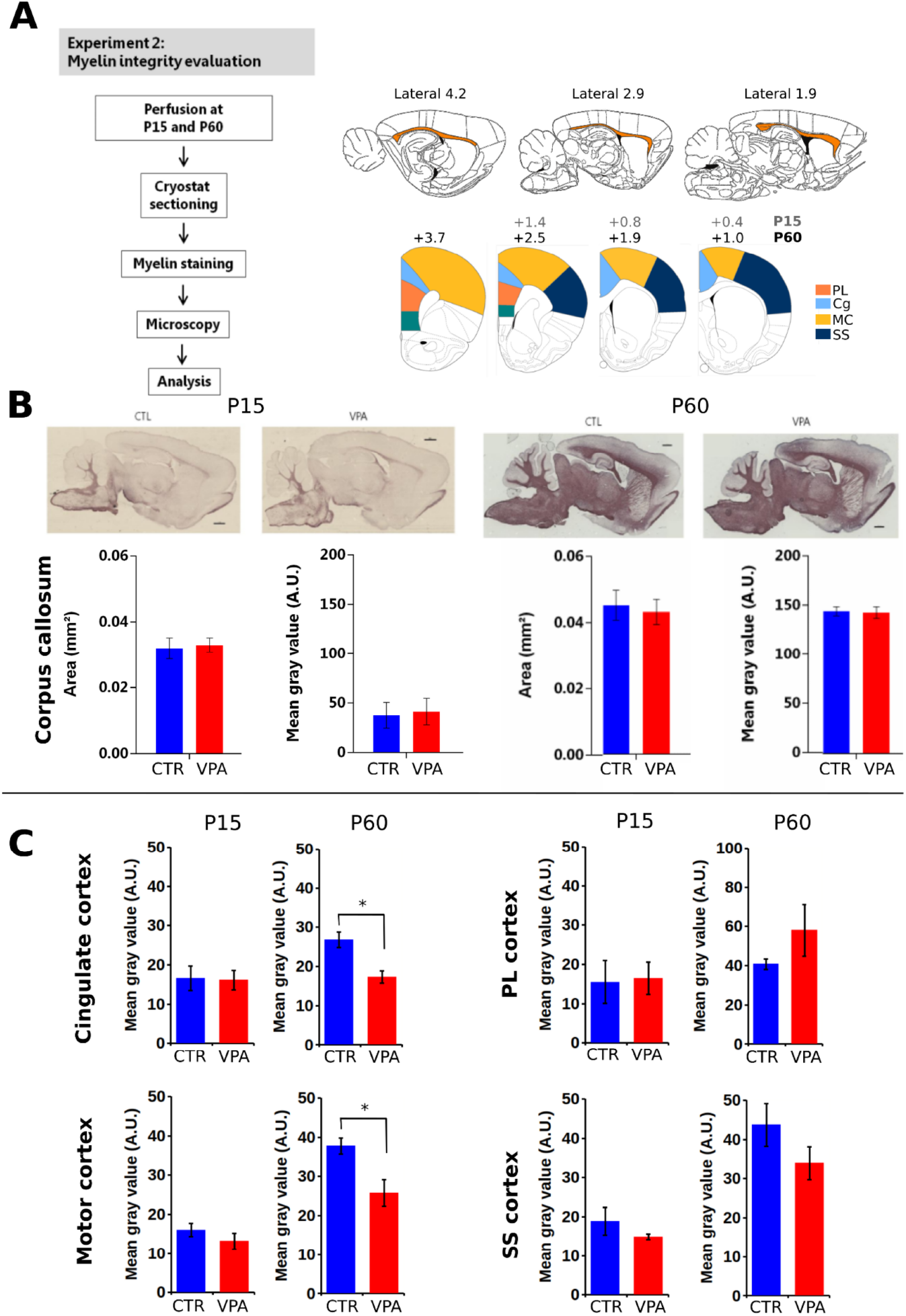

Analysis of the corpus callosum revealed no group differences in stained area or myelin intensity at either age (Fig. 5B). At P15, stained area (Mann–Whitney U, *p* = 0.65) and intensity (Mann–Whitney U, *p* = 0.48) were comparable between groups, and the same was observed at P60 (stained area: Mann–Whitney U, *p* = 0.99; intensity: Mann–Whitney U, *p* = 0.82). Analysis of the frontal cortex at P15, showed weak myelin staining consistent with previous reports (Mengler et al., 2014), and no group differences were observed across regions (Fig. 5B–C): total frontal cortex (Mann–Whitney U, *p* = 0.40), prelimbic cortex (Student’s t, *p* = 0.90), cingulate cortex (Mann–Whitney U, *p* = 0.85), motor cortex (Mann–Whitney U, *p* = 0.40), and somatosensory cortex (Mann–Whitney U, *p* = 0.62). However, at P60 robust myelin staining was evident, and VPA animals displayed significant reduction compared with controls in the cingulate cortex (Student’s t, *p* = 0.009; VPA: n = 4, 3 litters; CTL: n = 4, 2 litters) and motor cortex (Student’s t, *p* = 0.03; VPA: n = 4, 3 litters; CTL: n = 4, 2 litters). Trends to increased myelin staining in the prelimbic cortex and reduced myelin staining in the somatosensory cortex were observed, but the differences did not reach significance: prelimbic cortex (Mann– Whitney U, *p* > 0.05), or somatosensory cortex (Student’s t, *p* = 0.21).

### ALTERNATIVE SPLICING

Alternative splicing was analyzed for genes in which at least one isoform exhibited relative expression >75% of control levels and FPKM > 50. Four types of events were considered: exon skipping, alternative 3⍰ splice site, alternative 5⍰ splice site, and intron retention (Fig. 6). A total of 58 genes met these criteria: 17 were control-specific (29.5%), 24 were VPA-specific (41%), and 17 showed splicing in both groups. These genes produced 101 splicing variants, with 47 detected in controls (19 CTL-specific, 28 shared with VPA) and 54 in VPA animals (29 VPA-specific, 25 shared with controls).

**Fig. 6.**
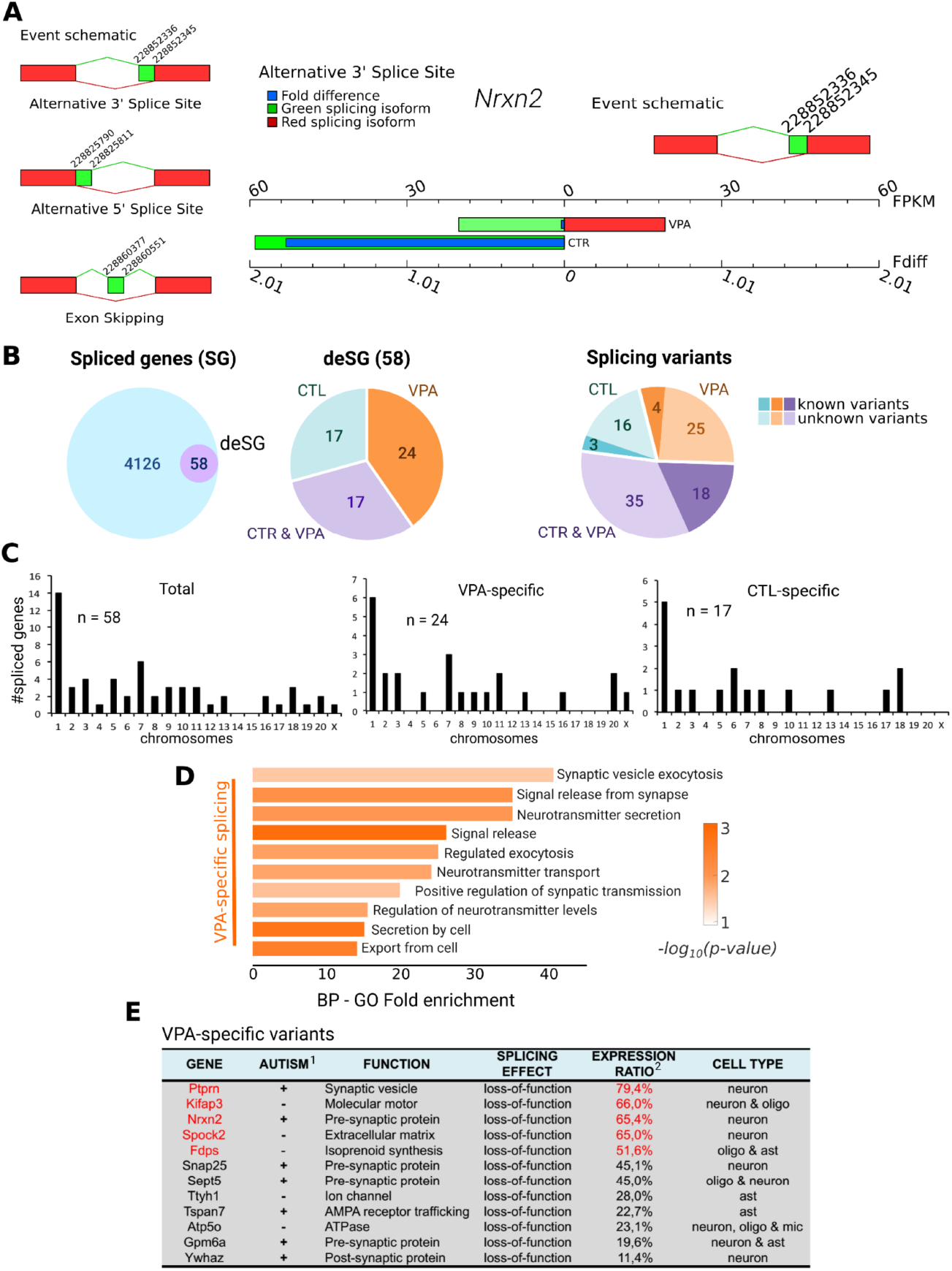

Variants were classified as known or novel based on the presence of RefSeq annotation. Among control-specific variants (n = 19), 84% (n = 16) were novel and 16% (n = 3) known. Among VPA-specific variants (n = 29), 86% (n = 25) were novel and 14% (n = 4) known. Overall, 25% (25/101) of all detected variants were annotated in RefSeq, whereas 75% (76/101) were novel transcripts (Fig. 6B).

Chromosomal mapping revealed a non-random distribution of spliced genes. VPA-specific splicing was enriched on chromosomes 9 (1 gene), 11 (2), 16 (1), 20 (2), and X (1), whereas control-specific splicing occurred on chromosomes 6 (2 genes), 17 (1), and 18 (2) (Fig. 6C).

Alternative splicing was detected in 29 of 103 downregulated genes (28%), comprising 17 exon-skipping, 6 alternative 31 splice-site, 6 alternative 5⍰ splice-site, and 8 intron-retention events (total = 37). Among upregulated genes, 33 of 76 (43%) exhibited splicing, including 30 exon-skipping, 19 alternative 3⍰, 15 alternative 51, and 7 intron-retention events (total = 71). In highly expressed genes (>30 FPKM), splicing was observed in 4 of 15 downregulated transcripts, including Enpp5, Zcchc12, and Synpr (>50 FPKM), and in 6 of 23 upregulated genes, with Sirt2, Acsbg1, Hbb, and Mobp exceeding 50 FPKM. In addition, poly-A variants were identified in nine genes (Olfm1, Cdc42, Cacng3, Sept5, Srsf3, Akr1a1, Brinp1, Gpm6b), while promoter variants were detected in 112 genes (data not shown).

VPA-specific variants were significantly enriched in synaptic regulation (Fig. 6D). Some of them showed to be non-functional splicing variants with significant relative expression levels (Fig. 6E). Together, these results demonstrate that autism-related to prenatal VPA exposure involves both changes in the frequency and chromosomal distribution of alternative splicing events, with a predominance of novel transcript variants, suggesting large-scale reorganization of synaptic and myelination-associated gene networks during early cortical development.

### 3.1 PLASMA BIOMARKER PROFILING

Serotonin has been proposed as a potential peripheral biomarker for autism. To investigate peripheral molecular signatures in early development, we first quantified plasma serotonin (5-HT) levels in VPA- and control-treated animals at P15 using HPLC with electrochemical detection. We then applied Fourier-transform infrared (FTIR) spectroscopy to test whether plasma spectral profiles provided reliable discrimination between VPA-treated and control animals (Fig. 7A-D).

**Fig. 7.**
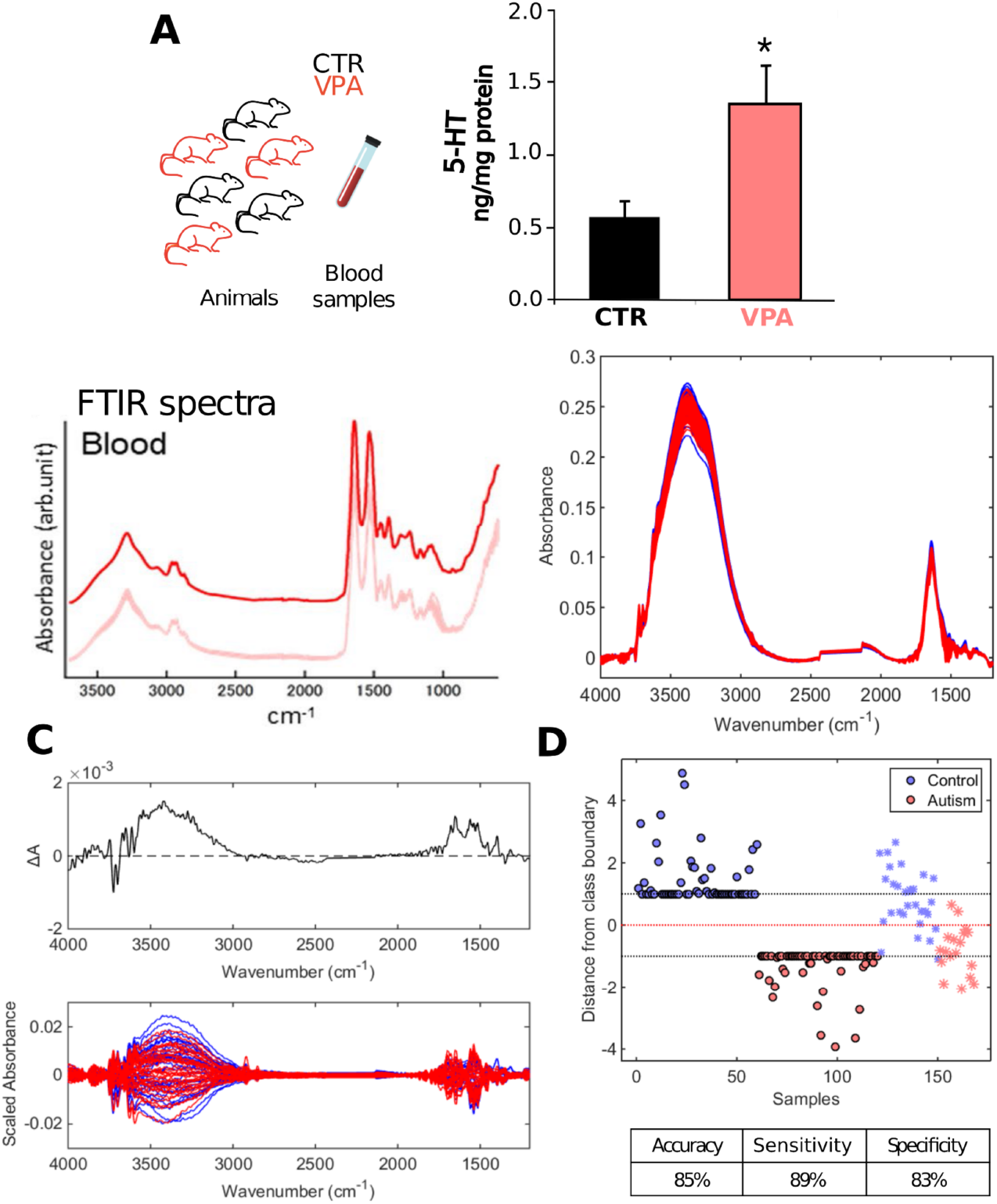

Our results showed that plasma 5-HT was significantly elevated in VPA animals compared with controls (+135%; p = 0.021, Student’s t-test). As for the FTIR analysis, pre-processed spectral data (Savitzky–Golay smoothing, window = 15 points, 2nd-order polynomial fitting; AWLS baseline correction) from CTL (n = 90) and VPA samples (n = 78) are shown in figure 7B. The spectral windows analyzed were 1178–2139 cm^−1^ and 2432–4000 cm^−1^, while the regions between 2139–2432 cm^−1^ (environmental CO_2_ interference) and 900–1178 cm^−1^ (lack of features) were excluded. The 1178– 2139 cm^−1^ region corresponds primarily to nucleic acids (PO_2_^−^ stretching in DNA/RNA, 1220–1240 cm^−1^), proteins (amide II, 1540–1550 cm^−1^; amide I, 1640–1660 cm^−1^), and lipids (CH_2_ bending, 1444– 1470 cm^−1^; C=O stretching, 1600–1800 cm^−1^; cis/trans C=C stretching, 1660–1670 cm^−1^), while the 2432–4000 cm^−1^ region includes lipids and fatty acids (CH_2_ stretching, 2850 cm^−1^; CH_3_ stretching, 2970 cm^−1^; CH stretching, 2800–3000 cm^−1^), and carbohydrates (O–H stretching, 3000–3700 cm^−1^) (Movasaghi et al., 2008).

Difference-between-means (DBM) and mean-centered spectra are shown in figure 7C. Positive signals (higher absorbance in controls) were evident at 3180–3500 cm^−1^ (O–H/N–H stretching in carbohydrates and proteins) and 1380–1700 cm^−1^ (CH_2_ bending in lipids/proteins and amide I/II bands), whereas VPA samples exhibited higher absorbance at 3685–3740 cm^−1^ (O–H/N–H stretching). Classification of samples into VPA and CTL groups was performed using a support vector machine (SVM) model trained with pre-processed spectral data (n = 60 CTL, n = 52 VPA). Model performance was optimized by venetian blinds cross-validation and tested on an independent set (n = 30 CTL, n = 26 VPA). The SVM achieved an accuracy of 85%, with sensitivity of 89% and specificity of 83%. Discriminant function analysis confirmed that all training samples were correctly classified, with the majority of test samples clustering within their true class boundaries (Fig. 7D).

## 4 DISCUSSION

Our findings reveal converging molecular and structural alterations of disrupted cortical development following prenatal valproic acid (VPA) exposure associated with autistic-like phenotypes in rodents. We demonstrated that prenatal VPA exposure triggers early-life alterations in frontal cortex gene expression —including synaptic gene upregulation (*Penk, C1ql3*) and myelin-related downregulation (*Mobp, PLP1*)—accompanied by abnormal alternative splicing events of synapse-associated genes. These molecular disruptions translated into adult deficits in cingulate and motor cortical myelination, elevated plasma serotonin, and distinctive serum FTIR spectral signatures capable of distinguishing VPA-exposed from control animals. Behaviorally, these neurobiological alterations coincide with autism-like phenotypes such as hyperlocomotion, repetitive grooming, and social indifference. Collectively, these findings indicate that prenatal VPA exposure disrupts coordinated developmental programs of synaptic and myelin gene regulation, leading to impaired cortical connectivity and altered behavioral outcomes, while also highlighting the potential of peripheral molecular signatures as translational biomarkers.

### Molecular Disruption of Synaptic- and Myelin-related genes in the Frontal Cortex

Transcriptomic profiling of the frontal cortex revealed a pronounced downregulation of myelin-associated genes and an upregulation of synaptic-related genes. These results are consistent with previous studies showing abnormal myelination, disruption of oligodendrocytic maturation and altered synaptic signaling following prenatal VPA exposure (Traetta et al., 2021; Liu et al., 2021; Uccelli et al., 2021.; Barrett et al., 2017; Go et al., 2012, PMID: 22101036; Kim et al., 2011). We also identified widespread VPA-specific alternative splicing, with a predominance of putatively non-functional isoforms enriched in synaptic transmission regulation. Given that VPA acts as a histone deacetylase (HDAC) inhibitor (Phiel et al., 2001), our results suggest that chromatin remodeling and splicing dysregulation may jointly contribute to transcriptomic instability leading to autism-like phenotype. Post-transcriptional regulation has recently gained attention as a mechanism shaping ASD susceptibility (Parikshak et al., 2016), and our data extend this notion by providing in vivo evidence of transcription and splicing disruption in the VPA model at early age (P15). Previous studies have shown changes in gene expression in the basolateral amygdala of VPA-treated rats at different developmental ages at P10, P21 and P35 (Barrett et al., 2017; Oguchi-Katayama et al. 2013).

### Myelination Deficits and Their Relevance to ASD Pathophysiology

There is growing evidence that white matter alterations contribute to ASD pathology by impairing neural communication efficiency across brain regions (Courchesne et al., 2001; Herbert et al., 2004; Ben Bashat et al., 2007; Roberts et al., 2013; Zhang et al., 2024). Diffusion tensor imaging (DTI) studies have reported lower fractional anisotropy and increased radial diffusivity in the white matter of ASD patients, suggesting axonal abnormalities and/or hypomyelination (Müller et al., 2011; Rane et al., 2015). Furthermore, electron microscopy analysis of the prefrontal cortex (PFC) of adult ASD patients revealed reduced myelin thickness in orbitofrontal cortex (OFC) axons, indicating disrupted connectivity in this region (Zikopoulos and Barbas, 2010). Transcriptomic studies have also highlighted myelin-related gene dysregulation in ASD. For instance, analysis of Brodmann area 19 (occipital cortex) and the cerebellum in ASD patients (2–60 years old) identified enrichment in myelin-associated genes (Ginsberg et al., 2012). Consistent with these findings, functional MRI studies have shown reduced long-range connectivity in ASD individuals during tasks requiring interregional coordination, further supporting the notion of impaired neural communication (Courchesne and Pierce, 2005; Just et al., 2007; Rane et al., 2015). Despite accumulating evidence linking white matter dysfunction to ASD, the underlying mechanisms remain poorly understood, particularly in ASD patients and animal models (Courchesne et al., 2001; Herbert et al., 2004; Ben Bashat et al., 2007; Pacey et al., 2013; Roberts et al., 2013; Wei et al., 2016).

Our histological analysis revealed significant reductions in myelin content in the cingulate and motor cortex of adult VPA-treated animals, while other frontal subregions and the corpus callosum were spared. This regional specificity suggests that molecular downregulation of myelin genes during early development (P15) translates into long-term, circuit-selective deficits at adulthood (P60). Thus, our findings reinforce the view that ASD is not characterized by uniform myelin disruption but by regionally selective alterations that may underlie impairments in social cognition and motor coordination. Previous studies have shown myelin dysfunctions later at juvenile stage (P36) of VPA-treated rats, which included reduced fraction of myelinated axons that were distorted and less compact in the corpus callosum in addition to intact myelination on the mPFC (Uccelli et al. 2021). Here, we similarly did not find abnormal organization of myelinated fibers in the mPFC (PL sub-region) and somatosensory cortex either at infancy or adulthood. In children with autism, it has been reported a similar pattern of lower levels of myelin-related proteins and elevated levels of synaptic proteins in the frontal cortex (Broek et a., 2014)

In other ASD animal models, myelination deficits have also been reported. The BTBR T+ltprtf/J mouse model exhibits reduced expression of myelin proteins, including *MBP, MAG*, and *PLP1* in cortical tissues (Wei et al., 2016). In the Fragile X syndrome (Fmr1 knockout) mouse model, it was observed a decreased expression of CNPase (an immature oligodendrocyte marker) and *MBP* (a mature oligodendrocyte marker) along with reduced myelin thickness and fewer myelinated axons in the cerebellum (Pacey et al., 2013). Different models of maternal immune activation (MIA) also show myelination abnormalities. In a prenatal influenza virus model (E16 exposure), *Mag* and *Mog* expression was decreased at birth (P0), upregulated at P14, and subsequently downregulated at P56 (Fatemi et al., 2009). Similarly, in a group B Streptococcus infection model, reduced white matter thickness, lower MBP density, and decreased *CC-1* staining were observed in the periventricular white matter (Bergeron et al., 2013). In the poly(I:C) viral mimetic model, myelin-related genes (*Mag, Mobp, Mal, Mog, Cldn11, Myrf*) and MOBP protein levels were downregulated in the medial PFC (Richetto et al., 2016; Zhang et al., 2020). Collectively, these findings reinforce the hypothesis that myelination impairments are a common pathological feature of ASD, with potential consequences for neural connectivity and function.

### FTIR Spectroscopy as a New Discriminating Tool for ASD

Peripheral analyses revealed elevated plasma serotonin levels in VPA-treated pups, in agreement with longstanding evidence of hyperserotonemia in a substantial subset of ASD patients (Anderson et al., 1987; Mulder et al., 2004). Beyond single-analyte measures, we applied Fourier-transform infrared (FTIR) spectroscopy coupled with machine learning analysis, achieving 85% accuracy in classifying VPA and control samples, highlighting its potential for future clinical applications. FTIR spectroscopy captures the vibrational profile of biomolecules in the plasma and has been explored as a potential diagnostic tool for ASD (Ogruc Ildiz et al., 2020) and proved effective in diagnosing neurodegenerative diseases such as Parkinson’s and Alzheimer’s (Ahmed et al., 2010; Paraskevaidi et al., 2017; Bury et al., 2019; Lilo et al., 2019). Considering that ASD diagnosis remains predominantly clinical, with no established biomolecular markers due to its multifactorial etiology, this is a novel finding, as spectroscopic approaches to ASD biomarker discovery remain underexplored. This approach find support in previous studies that have reported biochemical alterations in ASD individuals, including plasma hyperserotonemia, reduced endocannabinoid levels (2-AG and anandamide), and altered immune responses (cytokines and antibodies; Persico et al., 2002; Molloy et al., 2006; Karhson et al., 2018; Siniscalco et al., 2018). Our results provide proof-of-concept that peripheral molecular analyses are able to detect VPA-induced neurodevelopmental alterations. This underscores the potential of combining established biochemical markers with unbiased spectral profiling for translational biomarker pipelines.

## Conclusion

Here we provide evidence of how prenatal VPA exposure disrupts gene expression, alternative splicing, cortical myelination, and peripheral molecular profiles, culminating in long-term behavioral alterations. We observed developmental delays, hyperlocomotion, perseverative grooming, and impaired social preference—all consistent with prior reports in the model. By combining transcriptomics, histology, and spectroscopy with behavioral analysis, we demonstrate that early-life dysregulation of myelin and synaptic programs underlies both cortical and systemic changes with potential translational relevance. Importantly, the discovery of serum FTIR signatures that identify VPA animals highlights a minimally invasive avenue for biomarker development in ASD. Future studies should further dissect the causal role of splicing regulation in neurodevelopmental outcomes and validate spectroscopic biomarkers in clinical cohorts, paving the way toward integrative diagnostic tools for autism. Taken together, our findings strengthen the link between early molecular disruption and the emergence of enduring circuit and behavioral abnormalities in VPA-exposed animals.

## 5 DATA AVAILABILITY STATEMENT

The raw data supporting the conclusions of this article will be made available under request.

## 6 ETHICS STATEMENT

The animal study was reviewed and approved by the Ethical Committee for Animal Use in Research of the Federal University of Rio Grande do Norte (CEUA-UFRN, Protocol No. 013/2014).

## 7 AUTHOR CONTRIBUTIONS

C.A. - investigation, data curation, formal analysis, writing original draft

J.K. - software, data curation, formal analysis, writing review & editing

J.A.B. - investigation, data curation, formal analysis, visualization

R.S.B. - writing review & editing

S.R. - investigation, data curation, writing review & editing

C.L.M.M. - investigation, data curation, formal analysis, writing review & editing

M.C.D.S. - investigation, data curation, formal analysis, writing review & editing

K.M.G.L. - data curation, formal analysis, writing review & editing

S.S. - software, funding acquisition

R.N.R. - conceptualization, formal analysis, visualization, writing review & editing, resources, funding acquisition

## 8 FUNDING

This work was supported by CAPES (Coordenação de Aperfeiçoamento de Pessoal de Ensino Superior) providing funds for students and postdoc fellowships, CNPq (Conselho Nacional de Pesquisa) providing the research grants #430951/2016-7.

## 9 ACKNOWLEDGMENTS

The authors would like to thank the staff of the Brain Institute animal facility in the name of Mariana Campelo Medeiros and Josy Carolina Pontes for their assistance in maintaining the experimental animals.

## 10 CONFLICT OF INTEREST

The authors declare that the research was conducted in the absence of any commercial or financial relationships that could be construed as a potential conflict of interest.

## Notes

### Competing Interest Statement

The authors have declared no competing interest.

## REFERENCES

Ahmed, S. S. S. J., Santosh, W., Kumar, S., and Thanka Christlet, T. H. (2010). Neural network algorithm for the early detection of Parkinson’s disease from blood plasma by FTIR microspectroscopy. Vib. Spectrosc. 53, 181–188. doi:10.1016/j.vibspec.2010.01.019.

American Psychiatric Association (2013). Diagnostic and Statistical Manual of Mental Disorders. American Psychiatric Association doi:10.1176/appi.books.9780890425596.

Anders S and Huber W (2010). “Differential expression analysis for sequence count data.” Genome Biology, 11, pp. R106. 10.1186/gb-2010-11-10-r106.

Anderson, G. M. (2005). Serotonin in autism. In M. L. Bauman & T. L. Kemper (Eds.), The neurobiology of autism (2nd ed., pp. 303–318). Johns Hopkins University Press.

Anomal, R. F., de Villers-Sidani, E., Brandão, J. A., Diniz, R., Costa, M. R., and Romcy-Pereira, R. N. (2015). Impaired Processing in the Primary Auditory Cortex of an Animal Model of Autism. Front. Syst. Neurosci. 9. doi:10.3389/fnsys.2015.00158.

Ashburner, M., Ball, C.A., Blake, J.A., Botstein, D., Butler, H., Cherry, J.M., Davis, A.P., Dolinski, K., Dwight, S.S., Eppig, J.T., Harris, M.A., Hill, D.P., Issel-Tarver, L., Kasarskis, A., Lewis, S., Matese, J.C., Richardson, J.E., Ringwald, M., Rubin, G.M., Sherlock, G. Gene ontology: tool for the unification of biology. Nat Genet. May 2000;25(1):25–9

Ballabio, D., and Consonni, V. (2013). Classification tools in chemistry. Part 1: linear models. PLS-DA. Anal. Methods 5, 3790. doi:10.1039/c3ay40582f.

Barrett CE, Hennessey TM, Gordon KM, Ryan SJ, McNair ML, Ressler KJ, Rainnie DG. Developmental disruption of amygdala transcriptome and socioemotional behavior in rats exposed to valproic acid prenatally. Mol Autism. 2017 Aug 1;8:42. doi: 10.1186/s13229-017-0160-x. PMID: 28775827; PMCID: PMC5539636.

Ben Bashat, D., Kronfeld-Duenias, V., Zachor, D. A., Ekstein, P. M., Hendler, T., Tarrasch, R., et al. (2007). Accelerated maturation of white matter in young children with autism: A high b value DWI study. Neuroimage 37, 40–47. doi:10.1016/j.neuroimage.2007.04.060.

Bergeron, J. D. L., Deslauriers, J., Grignon, S., Fortier, L. C., Lepage, M., Stroh, T., et al. (2013). White Matter Injury and Autistic-Like Behavior Predominantly Affecting Male Rat Offspring Exposed to Group B Streptococcal Maternal Inflammation. Dev. Neurosci. 35, 504–515. doi:10.1159/000355656.

Bradford, M. M. (1976). A rapid and sensitive method for the quantitation of microgram quantities of protein utilizing the principle of protein-dye binding. Anal. Biochem. 72, 248–254. doi:10.1016/0003-2697(76)90527-3.

Broek, J., Guest, P., Rahmoune, H., & Bahn, S. (2014). Proteomic analysis of post mortem brain tissue from autism patients: evidence for opposite changes in prefrontal cortex and cerebellum in synaptic connectivity-related proteins. Molecular Autism, 5, 41–41. 10.1186/2040-2392-5-41.

Bromley, R. L., Mawer, G. E., Briggs, M., Cheyne, C., Clayton-Smith, J., García-Fiñana, M., et al. (2013). The prevalence of neurodevelopmental disorders in children prenatally exposed to antiepileptic drugs. J. Neurol. Neurosurg. & Psychiatry 84, 637LP–643. doi:10.1136/jnnp-2012-304270.

Bury, D., Morais, C. L. M., Paraskevaidi, M., Ashton, K. M., Dawson, T. P., and Martin, F. L. (2019). Spectral classification for diagnosis involving numerous pathologies in a complex clinical setting: A neuro-oncology example. Spectrochim. Acta Part A Mol. Biomol. Spectrosc. 206, 89–96. doi:10.1016/j.saa.2018.07.078.

Christensen, J., Grønborg, T. K., Sørensen, M. J., Schendel, D., Parner, E. T., Pedersen, L. H., et al. (2013). Prenatal valproate exposure and risk of autism spectrum disorders and childhood autism. JAMA 309, 1696–703. doi:10.1001/jama.2013.2270.

Christianson, A. L., Chesler, N., and Kromberg, J. G. (1994). Fetal valproate syndrome: clinical and neuro-developmental features in two sibling pairs. Dev. Med. Child Neurol. 36, 361–9. doi:10.1111/j.1469-8749.1994.tb11858.x.

Cortes, C., and Vapnik, V. (1995). Support-vector networks. Mach. Learn. 20, 273–297. doi:10.1007/BF00994018.

Courchesne, E., Karns, C. M., Davis, H. R., Ziccardi, R., Carper, R. A., Tigue, Z. D., et al. (2001). Unusual brain growth patterns in early life in patients with autistic disorder: An MRI study. Neurology 57, 245–254. doi:10.1212/WNL.57.2.245.

Courchesne, E., Mouton, P. R., Calhoun, M. E., Semendeferi, K., Ahrens-Barbeau, C., Hallet, M. J., et al. (2011). Neuron Number and Size in the Prefrontal Cortex of Children With Autism. JAMA 306, 2001. doi:10.1001/jama.2011.1638.

Courchesne, E., and Pierce, K. (2005). Brain overgrowth in autism during a critical time in development: implications for frontal pyramidal neuron and interneuron development and connectivity. Int. J. Dev. Neurosci. 23, 153–70. doi:10.1016/j.ijdevneu.2005.01.003.

Fatemi, S. H., Folsom, T. D., Reutiman, T. J., Abu-Odeh, D., Mori, S., Huang, H., et al. (2009). Abnormal expression of myelination genes and alterations in white matter fractional anisotropy following prenatal viral influenza infection at E16 in mice. Schizophr. Res. 112, 46–53. doi:10.1016/j.schres.2009.04.014.

Florea L, Song L, Salzberg SL. Thousands of exon skipping events differentiate among splicing patterns in sixteen human tissues. F1000Research 2012; 2.

Galante PAF, Sakabe NJ, Kirschbaum-Slager N, de Souza SJ. Detection and evaluation of intron retention events in the human transcriptome. RNA 2004; 10(5):757–765.

Galante PAF, Vidal DO, de Souza JE, Camargo AA, de Souza SJ. Sense-antisense pairs in mammals: functional and evolutionary considerations. Genome biology 2007; 8(3):R40.

Geschwind, D. H. (2011). Genetics of autism spectrum disorders. Trends Cogn. Sci. 15, 409–16. doi:10.1016/j.tics.2011.07.003.

Ginsberg, M. R., Rubin, R. A., Falcone, T., Ting, A. H., and Natowicz, M. R. (2012). Brain Transcriptional and Epigenetic Associations with Autism. PLoS One 7, e44736. doi:10.1371/journal.pone.0044736.

Go, H. S., Kim, K. C., Choi, C. S., Jeon, S. J., Kwon, K. J., Han, S.-H., et al. (2012). Prenatal exposure to valproic acid increases the neural progenitor cell pool and induces macrocephaly in the rat brain via a mechanism involving the GSK-3β/β-catenin pathway. Neuropharmacology 63, 1028–1041. doi:10.1016/j.neuropharm.2012.07.028.

Gogolla, N., LeBlanc, J. J., Quast, K. B., Südhof, T. C., Fagiolini, M., and Hensch, T. K. (2009). Common circuit defect of excitatory-inhibitory balance in mouse models of autism. J. Neurodev. Disord. 1, 172–181. doi:10.1007/s11689-009-9023-x.

Hara, Y., Maeda, Y., Kataoka, S., Ago, Y., Takuma, K., and Matsuda, T. (2012). Effect of Prenatal Valproic Acid Exposure on Cortical Morphology in Female Mice. J. Pharmacol. Sci. 118, 543–546. doi:10.1254/jphs.12025SC.

Herbert, M. R., Ziegler, D. A., Makris, N., Filipek, P. A., Kemper, T. L., Normandin, J. J., et al. (2004). Localization of white matter volume increase in autism and developmental language disorder. Ann. Neurol. 55, 530–540. doi:10.1002/ana.20032.

HTSeq — A Python framework to work with high-throughput sequencing data Bioinformatics (2014), in print, online at doi:10.1093/bioinformatics/btu638.

Ingram, J. L., Peckham, S. M., Tisdale, B., and Rodier, P. M. (2000). Prenatal exposure of rats to valproic acid reproduces the cerebellar anomalies associated with autism. Neurotoxicol. Teratol. 22, 319–24. doi:10.1016/s0892-0362(99)00083-5.

Just, M. A., Cherkassky, V. L., Keller, T. A., Kana, R. K., and Minshew, N. J. (2007). Functional and Anatomical Cortical Underconnectivity in Autism: Evidence from an fMRI Study of an Executive Function Task and Corpus Callosum Morphometry. Cereb. Cortex 17, 951–961. doi:10.1093/cercor/bhl006.

Karhson, D. S., Krasinska, K. M., Dallaire, J. A., Libove, R. A., Phillips, J. M., Chien, A. S., et al. (2018). Plasma anandamide concentrations are lower in children with autism spectrum disorder. Mol. Autism 9, 18. doi:10.1186/s13229-018-0203-y.

Khazipov, R., Zaynutdinova, D., Ogievetsky, E., Valeeva, G., Mitrukhina, O., Manent, J.-B., et al. (2015). Atlas of the Postnatal Rat Brain in Stereotaxic Coordinates. Front. Neuroanat. 9. doi:10.3389/fnana.2015.00161.

Kim, K. C., Kim, P., Go, H. S., Choi, C. S., Yang, S.-I., Cheong, J. H., et al. (2011). The critical period of valproate exposure to induce autistic symptoms in Sprague–Dawley rats. Toxicol. Lett. 201, 137–142. doi:10.1016/j.toxlet.2010.12.018.

Kroll JE, Galante PA, Ohara DT, Navarro FC, Ohno-Machado L, de Souza SJ. SPLOOCE: a new portal for the analysis of human splicing variants. RNA biology 2012; 9(11):1339–1343.

Kroll JE, Kim J, Ohno-Machado L, de Souza SJ. (2015) Splicing Express: a software suite for alternative splicing analysis using next-generation sequencing data. PeerJ 3:e1419 10.7717/peerj.1419

Lajeunie, E., Barcik, U., Thorne, J. A., Ghouzzi, V. El, Bourgeois, M., and Renier, D. (2001). Craniosynostosis and fetal exposure to sodium valproate. J. Neurosurg. 95, 778–782. doi:10.3171/jns.2001.95.5.0778.

Lilo, T., Morais, C., Ashton, K., Pardilho, A., Dawson, T., Gurusinghe, N., et al. (2019). Predicting meningioma recurrence using spectrochemical analysis of tissues and subsequent predictive computational algorithms. Neuro. Oncol. 21, iv5–iv5. doi:10.1093/neuonc/noz167.020.

Liu H, Tan M, Cheng B, Wang S, Xiao L, Zhu J, Wu Q, Lai X, Zhang Q, Chen J, Li T. Valproic Acid Induces Autism-Like Synaptic and Behavioral Deficits by Disrupting Histone Acetylation of Prefrontal Cortex ALDH1A1 in Rats. Front Neurosci. 2021 Apr 28;15:641284. doi: 10.3389/fnins.2021.641284. PMID: 33994921; PMCID: PMC8113628.

Livak, K. J., and Schmittgen, T. D. (2001). Analysis of Relative Gene Expression Data Using Real-Time Quantitative PCR and the 2−ΔΔCT Method. Methods 25, 402–408. doi:10.1006/meth.2001.1262.

Mengler, L., Khmelinskii A., Diedenhofen M., Po C., Staring M., Lelieveldt B., Hoehn, M. (2014). Brain maturation of the adolescent rat cortex and striatum: changes in volume and myelination. NeuroImage, 84:35–44.

Molloy, C. A., Morrow, A. L., Meinzen-Derr, J., Schleifer, K., Dienger, K., Manning-Courtney, P., et al. (2006). Elevated cytokine levels in children with autism spectrum disorder. J. Neuroimmunol. 172, 198–205. doi:10.1016/j.jneuroim.2005.11.007.

Moore, S. J., Turnpenny, P., Quinn, A., Glover, S., Lloyd, D. J., Montgomery, T., et al. (2000). A clinical study of 57 children with fetal anticonvulsant syndromes. J. Med. Genet. 37, 489–97. doi:10.1136/jmg.37.7.489.

Morais, C. L. M., Costa, F. S. L., and Lima, K. M. G. (2017). Variable selection with a support vector machine for discriminating Cryptococcus fungal species based on ATR-FTIR spectroscopy. Anal. Methods 9, 2964–2970. doi:10.1039/C7AY00428A.

Morais, C. L. M., Paraskevaidi, M., Cui, L., Fullwood, N. J., Isabelle, M., Lima, K. M. G., et al. (2019a). Standardization of complex biologically derived spectrochemical datasets. Nat. Protoc. 14, 1546–1577. doi:10.1038/s41596-019-0150-x.

Morais, C. L. M., Santos, M. C. D., Lima, K. M. G., and Martin, F. L. (2019b). Improving data splitting for classification applications in spectrochemical analyses employing a random-mutation Kennard-Stone algorithm approach. Bioinformatics 35, 5257–5263. doi:10.1093/bioinformatics/btz421.

Movasaghi, Z., Rehman, S., and ur Rehman, D. I. (2008). Fourier Transform Infrared (FTIR) Spectroscopy of Biological Tissues. Appl. Spectrosc. Rev. 43, 134–179. doi:10.1080/05704920701829043.

Müller, R.-A., Shih, P., Keehn, B., Deyoe, J. R., Leyden, K. M., and Shukla, D. K. (2011). Underconnected, but How? A Survey of Functional Connectivity MRI Studies in Autism Spectrum Disorders. Cereb. Cortex 21, 2233–2243. doi:10.1093/cercor/bhq296.

Ogruc Ildiz, G., Bayari, S., Karadag, A., Kaygisiz, E., and Fausto, R. (2020). Fourier Transform Infrared Spectroscopy Based Complementary Diagnosis Tool for Autism Spectrum Disorder in Children and Adolescents. Molecules 25. doi:10.3390/molecules25092079.

Pacey, L. K. K., Xuan, I. C. Y., Guan, S., Sussman, D., Henkelman, R. M., Chen, Y., et al. (2013). Delayed myelination in a mouse model of fragile X syndrome. Hum. Mol. Genet. 22, 3920–3930. doi:10.1093/hmg/ddt246.

Paraskevaidi, M., Morais, C. L. M., Lima, K. M. G., Snowden, J. S., Saxon, J. A., Richardson, A. M. T., et al. (2017). Differential diagnosis of Alzheimer’s disease using spectrochemical analysis of blood. Proc. Natl. Acad. Sci. 114, E7929–E7938. doi:10.1073/pnas.1701517114.

Pardo, C. A., and Eberhart, C. G. (2007). The Neurobiology of Autism. Brain Pathol. 17, 434–447. doi:10.1111/j.1750-3639.2007.00102.x.

Paxinos, G., and Watson, C. (2008). The Rat Brain in Stereotaxic Coordinates. 6th ed. Elsevier.

Persico, A. M., Pascucci, T., Puglisi-Allegra, S., Militerni, R., Bravaccio, C., Schneider, C., et al. (2002). Serotonin transporter gene promoter variants do not explain the hyperserotonemia in autistic children. Mol. Psychiatry 7, 795–800. doi:10.1038/sj.mp.4001069.

Phiel, C. J., Zhang, F., Huang, E. Y., Guenther, M. G., Lazar, M. A., and Klein, P. S. (2001). Histone Deacetylase Is a Direct Target of Valproic Acid, a Potent Anticonvulsivant has been explored as a potential diagnostic tool for ASD J. Biol. Chem. 276, 36734–36741. doi:10.1074/jbc.M101287200.

Rane, P., Cochran, D., Hodge, S. M., Haselgrove, C., Kennedy, D. N., and Frazier, J. A. (2015). Connectivity in Autism. Harv. Rev. Psychiatry 23, 223–244. doi:10.1097/HRP.0000000000000072.

Richetto, J., Chesters, R., Cattaneo, A., Labouesse, M. A., Gutierrez, A. M. C., Wood, T. C., et al. (2016). Genome-Wide Transcriptional Profiling and Structural Magnetic Resonance Imaging in the Maternal Immune Activation Model of Neurodevelopmental Disorders. Cereb. Cortex. doi:10.1093/cercor/bhw320.

Roberts A, Pimentel H, Trapnell C, Pachter L. Identification of novel transcripts in annotated genomes using RNA-Seq. Bioinformatics 2011; 27(17):2325–2329.

Roberts, T. P. L., Lanza, M. R., Dell, J., Qasmieh, S., Hines, K., Blaskey, L., et al. (2013). Maturational differences in thalamocortical white matter microstructure and auditory evoked response latencies in autism spectrum disorders. Brain Res. 1537, 79–85. doi:10.1016/j.brainres.2013.09.011.

Sammeth M, Foissac S, Guigó R. A general definition and nomenclature for alternative splicing events. PLoS Comput Biol 2008; 4:e1000147.

Schindelin, J., Arganda-Carreras, I., Frise, E., Kaynig, V., Longair, M., Pietzsch, T., … Cardona, A. (2012). Fiji: an open-source platform for biological-image analysis. Nature Methods, 9(7), 676–682. doi:10.1038/nmeth.2019

Schmued, L., Bowyer, J., Cozart, M., Heard, D., Binienda, Z., and Paule, M. (2008). Introducing Black-Gold II, a highly soluble gold phosphate complex with several unique advantages for the histochemical localization of myelin. Brain Res. 1229, 210–7. doi:10.1016/j.brainres.2008.06.129.

Schneider, T., and Przewłocki, R. (2005). Behavioral alterations in rats prenatally exposed to valproic acid: animal model of autism. Neuropsychopharmacology 30, 80–9. doi:10.1038/sj.npp.1300518.

Schneider, T., Roman, A., Basta-Kaim, A., Kubera, M., Budziszewska, B., Schneider, K., et al. (2008). Gender-specific behavioral and immunological alterations in an animal model of autism induced by prenatal exposure to valproic acid. Psychoneuroendocrinology 33, 728–740. doi:10.1016/j.psyneuen.2008.02.011.

Siniscalco, D., Schultz, S., Brigida, A., and Antonucci, N. (2018). Inflammation and Neuro-Immune Dysregulations in Autism Spectrum Disorders. Pharmaceuticals 11, 56. doi:10.3390/ph11020056.

Skefos, J., Cummings, C., Enzer, K., Holiday, J., Weed, K., Levy, E., et al. (2014). Regional Alterations in Purkinje Cell Density in Patients with Autism. PLoS One 9, e81255. doi:10.1371/journal.pone.0081255.

Trapnell, C., Pachter, L., and Salzberg, S. L. (2009). TopHat: discovering splice junctions with RNA-Seq. Bioinformatics 25, 1105–1111. doi:10.1093/bioinformatics/btp120.

Uccelli NA, Codagnone MG, Traetta ME, Levanovich N, Rosato Siri MV, Urrutia L, Falasco G, Vázquez S, Pasquini JM, Reinés AG. Neurobiological substrates underlying corpus callosum hypoconnectivity and brain metabolic patterns in the valproic acid rat model of autism spectrum disorder. J Neurochem. 2021 Oct;159(1):128–144. doi: 10.1111/jnc.15444. Epub 2021 Jun 28. PMID: 34081798.

Vedi, M., Smith, J.R., Hayman, G.T., Tutaj, M., Brodie, K.C., De Pons, J.L., Demos, W.M., Gibson, A.C., Kaldunski, M.L., Lamers, L., Laulederkind, S.J.F., Thota, J., Thorat, K., Tutaj, M.A., Wang, S, Zacher, S., Dwinell, M.R., Kwitek, A.E. 2022 updates to the Rat Genome Database: a Findable, Accessible, Interoperable, and Reusable (FAIR) resource, Genetics, Volume 224, Issue 1, May 2023, iyad042, 10.1093/genetics/iyad042.

Wei, H., Ma, Y., Liu, J., Ding, C., Hu, F., and Yu, L. (2016). Proteomic analysis of cortical brain tissue from the BTBR mouse model of autism: Evidence for changes in STOP and myelin-related proteins. Neuroscience 312, 26–34. doi:10.1016/j.neuroscience.2015.11.003.

Yu G, Wang L, Han Y and He Q (2012). “clusterProfiler: an R package for comparing biological themes among gene clusters.” OMICS: A Journal of Integrative Biology, 16(5), pp. 284–287. 10.1089/omi.2011.0118.

Zhang, S., Jiang, L., Hu, Z., Liu, W., Yu, H., Chu, Y., Wang, J., & Chen, Y. (2024). T1w/T2w ratio maps identify children with autism spectrum disorder and the relationships between myelin-related changes and symptoms. Progress in Neuro-Psychopharmacology and Biological Psychiatry, 134.

Zhang X-F, Chen T, Yan A, Xiao J, Xie Y-L, Yuan J, Chen P, Wong AO-L, Zhang Y and Wong N-K (2020) Poly(I:C) Challenge Alters Brain Expression of Oligodendroglia-Related Genes of Adult Progeny in a Mouse Model of Maternal Immune Activation. Front. Mol. Neurosci. 13:115. doi: 10.3389/fnmol.2020.00115

Zikopoulos, B., and Barbas, H. (2010). Changes in Prefrontal Axons May Disrupt the Network in Autism. J. Neurosci. 30, 14595–14609. doi:10.1523/JNEUROSCI.2257-10.2010.

