## Supplementary material for "Developmental Dysregulation of Synaptic and Myelin-Related Genes in Frontal Cortex and Serum Infrared Spectroscopy Signature in the Valproic Acid Model of Autism": Figure_Legends

**Figure 1:** **A,** Experimental design showing the generation of the rodent model of autism, and the timing of the behavioral tests and tissue collection. Pregnant females received a systemic injection of valproic acid (500 mg/Kg i.p.; on E12.5) and postnatal development of pups was monitored until P35. Biological samples were collected from P8 to P60, spanning infancy, adolescence and adulthood for tissue and molecular analysis. **B**, VPA-treated females showed similar litter size as compared to saline-treated controls. **C-H,** Animals prenatally exposed to VPA show autistic-like developmental and behavioral deficits. **C,** Newborns exposed to VPA showed a delay in eye opening, a hallmark of postnatal development. **D**, Control and VPA-treated rats showed similar gains of body weight between P7 and P21. However, after weaning (P28-P35), VPA-treated animals showed reduced weight gain compared to controls. **E**, VPA-treated rats showed increased locomotor activity. **F-G,** longer grooming episodes. **H,** and social interaction deficit. *, p < 0.05; **, p < 0.01 and ***, p < 0.001 (Student t-test or Mann-Whitney U test).

**Figure 2**: RNA sequencing profile of the frontal cortex of P15 newborn rats (VPA-treated vs. age-matched controls). **A,** Tissue dissection and basic RNAseq statistics. **B**, Volcano plot of differentially cortical expression between VPA and controls. **C**, Gene expression plot in FPKM units highlighting the down-regulation of myelin-related genes (Plp1, Mobp, Cnp, Cldn11, Mag: *in blue*) in VPA-treated rats compared to controls. **D**, Gene ontology analysis shows a significant enrichment of GABAergic, dopaminergic, glutamatergic and neuropeptide signaling among up-regulated genes. In contrast, myelin-related genes were significantly more represented among down-regulated genes. **E**, KEGG pathways analysis shows significant enrichment in neuroactive ligand-receptor interaction, calcium-signaling pathway and axon guidance. **F,** Differentially expressed genes in VPA-treated animals have 19 common genes present in the SFARI gene database (Q2-2025 Report), which corresponds to ~10% of the VPA-related DEGs.

**Figure 3**: Distribution of autism-related genes in human and rat chromosomes. **A**, Enrichment of autism-related genes across human chromosomes. In *circles*, the expected number of genes to be affected, considering the fraction of protein-coding genes in each chromosome and the *bars*, the actual number of autism-related SFARI score 1-2 genes reported (SFARI database, Q2-2025 Report). *Dotted-circles* indicate chromosomes enriched for autism-related genes. **B,** Human-to-rat chromosome mapping of enriched SFARI score 1-2 genes based on the conserved synteny mapping between the two genomes - humanGRCh38 (primary) and rat Rnor_6.0 assemblies/RGD-VCMap v1.0.3. **C,** Enrichment of VPA-modulated gene expression across the rat chromosomes. In *circles*, the expected number of genes to be affected, considering the fraction of protein-coding in each chromosome and the *bars*, the actual number of DEGs induced by prenatal exposure to VPA. *Dotted-circles* indicate chromosomes enriched for VPA-modulated genes. **D,** Number of up-regulated (*dark green*) and down-regulated (*light green*) genes induced by prenatal exposure to VPA and their location across the rat chromosomes. **E,** Fraction of VPA-induced DEGs in each chromosome, i.e. number of differentially expressed genes in the chromosome divided by the total number of genes in that same chromosome.

**Figure 4:** Validation of the cortical differential expression between P15 newborn VPA-treated rats and controls. **A**, Experimental flowchart. Frontal cortex samples were identically collected and prepared for qPCR analysis. **B-C**, VPA-treated animals exhibited significantly lower frontal cortex expression of the myelin-associated genes Mobp and Plp1 than controls. **C**, VPA-treated animals exhibited significantly higher frontal cortex expression of the synaptic transmission-associated genes Penk and C1ql3 than controls. *, p < 0.05 (Student t-test).

**Figure 5:** Altered myelin development in VPA-treated animals. **A,** Histological procedure to quantify brain tissue myelin. Brain regions analyzed are depicted in colors. Corpus callosum (in *orange*, *top*). Motor cortex (MC, *yellow*), Cingulate cortex (Cg, *blue*), Prelimbic cortex (PL, *orange*), somatosensory cortex (SS, *dark blue*) **B,** Corpus callosum myelin staining in brain sections from VPA-treated and control animals at ages P15 and P60. **C,** Significant lower myelin levels were observed in the Cg and MC of VPA-treated animals as compared to controls. *, p < 0.05 (Student t-test)

**Figure 6:** Alternative splicing in the frontal cortex of VPA-treated rats. **A,** Splicing events evaluated: 3’ splice site, 5’ splice site, exon skipping and intron retention (*not shown*). Example of the *nrxn2* gene in control and VPA-treated samples. ; **B,** Differential expression in alternative spliced genes (deSG). Fifty-eight genes had a differential expression of some splicing variant in VPA-treated animals compared to controls. Out of the 58 deSG, 24 genes occurred specifically in the VPA model of autism and 17 were shared with controls. Unknown splicing variants represented the majority of differentially expressed variants in VPA-treated animals. **C,** Chromosomal distribution of alternatively spliced genes. **D,** Gene ontology analysis of VPA-specific deSG shows a significant enrichment of genes related to synaptic transmission. **E,** List of VPA-specific splicing variants with predicted loss of function - some of them previously associated with *autism* based on the SFARI database^1^. *Expression* is shown as ratios relative to the summed counts of all variants^2^. *Cell type* indicates the cell population in which relative expression is enriched.

**Figure 7:** Serotonin levels and FTIR spectroscopy of blood plasma from VPA-treated animals. **A,** Significantly higher levels of serotonin were observed in experimental animals compared to controls. *, p < 0.05 (Student t-test). **B,** FTIR spectra. Full spectrum example and individual superposed spectra after removal of CO_2_ interference and absorptions below 900 cm^-1^ region (no biological information). The region between 900–1177 cm^-1^ was also removed since no absorption band was observed. Savitzky-Golay smoothing (window = 15 points, 2nd order polynomial), automatic weighted least squares baseline correction (3rd order polynomial fitting). **C,** Difference-between-mean support vectors spectra and mean-centred spectra (*top*). Higher influence for controls at: 3500 – 3180 cm^-1^ and 1700 – 1380 cm^-1^ and higher influence for experimental (VPA) samples at: 3740 – 3685 cm^-1^. Individual scaled spectra at the bottom: VPA-treated (*red*) and control samples (*blue*) **D,** Support Vector Machine classification of blood FTIR spectra from VPA-treated and controls.
